# OneOPES, a combined enhanced sampling method to rule them all

**DOI:** 10.1101/2023.03.06.531337

**Authors:** Valerio Rizzi, Simone Aureli, Narjes Ansari, Francesco Luigi Gervasio

## Abstract

Enhanced sampling techniques have revolutionised molecular dynamics (MD) simulations, enabling the study of rare events and the calculation of free energy differences in complex systems. One of the main families of enhanced sampling techniques uses physical degrees of freedom called collective variables (CVs) to accelerate a system’s dynamics and recover the original system’s statistics. However, encoding all the relevant degrees of freedom in a limited number of CVs is challenging, particularly in large biophysical systems. Another category of techniques, such as parallel tempering, simulates multiple replicas of the system in parallel, without requiring CVs. However, these methods may explore less relevant high-energy portions of the phase space and become computationally expensive for large systems. To overcome the limitations of both approaches, we propose a replica exchange method called OneOPES that combines the power of multi-replica simulations and CV-based enhanced sampling. This method efficiently accelerates the phase space sampling without the need for ideal CVs, extensive parameters fine tuning nor the use of a large number of replicas, as demonstrated by its successful applications to protein-ligand binding and protein folding benchmark systems. Our approach shows promise as a new direction in the development of enhanced sampling techniques for molecular dynamics simulations, providing an efficient and robust framework for the study of complex and unexplored problems.

**Table of Content Graphic:** 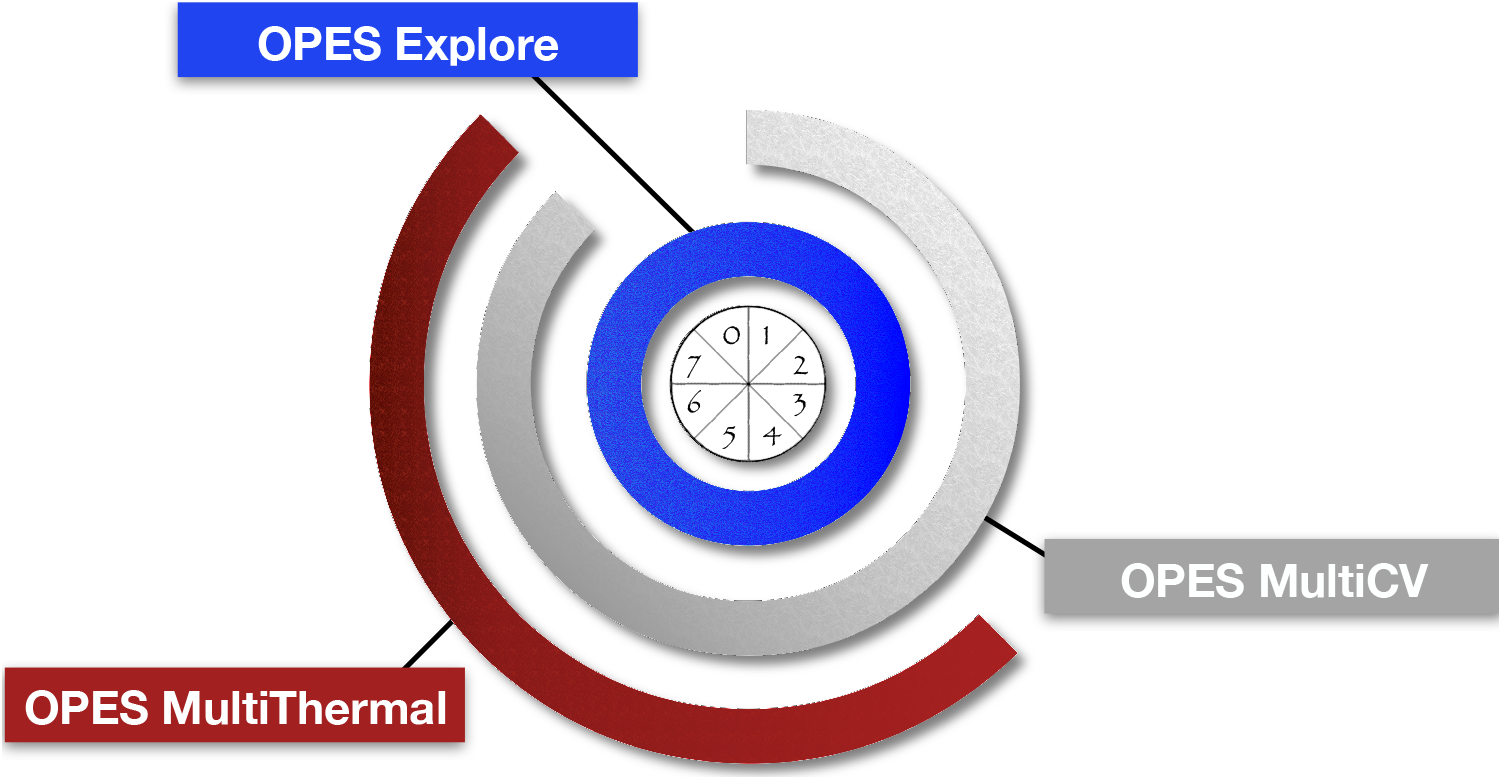

## Introduction

The development of enhanced sampling techniques has allowed molecular dynamics (MD) simulations to explore considerably longer time-scales, unlocking the possibility of routinely studying rare events and calculating free energy differences in a number of complex problems^1–4^.

One of the main families of enhanced sampling techniques tackles the problem by accelerating a system’s dynamics along selected physical degrees of freedom (DOFs) called collective variables (CVs) and by recovering the original system’s statistics by factoring out the deposited bias. In time, many CV-based sampling methods have been developed that are actively used by the community^4^. Typically, the maximum number of CVs that can be simultaneously biased is limited, due to the exponentially increasing cost of exhaustively exploring multi-dimensional free energy spaces. The precursor of such methods is Umbrella Sampling^5^, while Metadynamics^6^ is one of the most popular. In Well-Tempered Metadynamics (WT-MetaD)^7^, one iteratively builds a bias potential that, for a long enough time, is proven to converge and provide exact statistical properties^8^. However, the definition of “long enough time” is determined by the CVs’ capability to capture all the slow DOFs of a system that are relevant for the process studied.

The quality of the CVs crucially determines the simulation timescale needed to reach convergence^9^, as the slowest ignored DOFs dictate the speed at which transitions between metastable states occur, making the standard MD timescale problem reappear and translate into a search for better CVs. Encoding all the relevant DOFs in a limited number of CVs is no easy task and it is especially unfeasible in the large and complex systems that characterise the biophysical world. Common CVs such as distances and angles are quite straightforward to figure out and do not require extensive knowledge of the problem at hand, but they are hopeless to capture complex many-body transitions. More sophisticated techniques to build optimal CVs by combining simpler ones, or by finding an optimal path between initial and final states, or by using machine learning on large datasets have been successful, but they still often require significant human effort or large amounts of data^10–29^.

Another category of techniques addresses the timescale problem through parallelism by simulating multiple realisations of the system, or replicas, at the same time^30–32^. Typically, these methods do not require the introduction of CVs to be biased. Among them, replica exchange methods simulate parallel replicas differing from each other through the variation of internal system’s parameters^30,33,34^ and periodically attempt exchanging their coordinates, given a physical acceptance criterion. One prominent example of such methods is Parallel tempering^33,35^ and its variants^36,37^, where a progressive temperature increase in the replicas leads to all kinetic barriers being lowered and enthalpy-driven processes being accelerated. Parallel tempering methods have the advantage of being able to sample large phase space regions without the necessity to reach very long timescales and without being limited by missing slow DOFs in the CVs definition. However, high temperatures and no CV-defined direction can lead the simulations to explore less relevant high-energy portions of the phase space, reducing overall efficiency and not improving their capability to overcome entropic barriers^38,39^. Furthermore, in large systems, a considerable number of replicas must often be employed to ensure effective exchanges, making Parallel Tempering methods expensive^40^. While solutions to some of these limitations are arising^41,42^, a possible way forward is to combine the power of multi-replica simulations and CV-based enhanced sampling.

A number of successful methods have attempted, to varying degrees, to combine the power of both approaches, including Multiple-Walkers Metadynamics (MW-MetaD)^43^, Parallel-Tempering Metadynamics (PT-MetaD)^44,45^, Bias-Exchange Metadynamics (BE-MetaD)^46^ and further evolutions^47–53^. Many of these methods showed significant promise as the combination of CV and replica-based algorithms is able to efficiently accelerate the crossing of both enthalpic and entropic barriers without the necessity of an optimal CV development, as shown by their successful application to complex biological systems^54,55^. However, they also inherited some of the intrinsic limitations of their predecessors, namely a reduced but still present dependence of the free energy convergence on the quality of the CVs, a problematic setup of optimal parameters, and a significant computational cost. The rise of novel enhanced sampling techniques such as On-the-fly Probability Enhanced Sampling (OPES)^56^ and its variants^57,58^ has prompted us to formulate an OPES-based replica exchange method. Its aim is to provide a framework that produces converged results at a reasonable computational cost while being less reliant on the setup parameters and the CV quality. The overall strategy exploits the qualities of a combination of existing OPES variants in a parallel strategy that we call *OneOPES*.

In standard OPES^56^, one estimates the unbiased probability distribution by depositing weighted Gaussian kernels along chosen CVs. In a thought-provoking paper^58^, it was shown that, when CVs are excellent, the rapidly converging bias potential leads to a high transition frequency and very accurate results in a short computational time. On the other hand, when CVs are suboptimal, the OPES bias potential determines a slow phase space exploration that in turn forces simulations to extend for long times before reaching convergence. OPES Explore^58^ addresses this point and builds a more rapidly varying bias potential which leads to a faster phase space exploration, at the price of a slower and noisier convergence. Meanwhile, another conceptually different OPES variant called OPES MultiThermal^57^ has been developed, where a system simultaneously visits a range of temperature distributions without changing the thermostat or having to run multiple replicas.

In OneOPES, inspired by the approach of Ref. 48, we set up a mixture of replicas including OPES variants of different exploratory intensities and combine them with explicit replica exchange. All replicas include one OPES Explore bias that sets a common baseline by building a bias potential over a set of leading CVs. The first replica is the most convergence-focused replica and only includes this standard OPES Explore bias potential. As such, it will be used to calculate equilibrium properties through reweighting. Higher-order replicas are more exploratory and may include OPES Explore biases applied on a number of other CVs. These extra biases are weaker and updated more infrequently than the leading bias. Their purpose is to complement the leading bias by accelerating the sampling of transversal DOFs that are not included in the leading CVs, as done in Ref. 46,48. The most exploratory replicas also include OPES MultiThermal so that the effect of suboptimal CVs is further mitigated and all kinetic barriers are lowered. In a nutshell, exchanges between exploratory and convergence-dedicated replicas ensure that the former simulations bring variety into the latter ones, as convergence-dedicated ones moderate the *exuberance* of the exploratory ones.

We use OneOPES in combination with standard but suboptimal CVs that would undermine the convergence of the reconstructed free energy when used in combination with standard CV-based approaches and test it on a set of case studies that presents a diverse set of difficulties and requirements. As a stringent convergence criterion, for each system we perform a set of five completely independent simulations and evaluate their average outcome. At first, we simulate a system that is commonly used in enhanced sampling algorithm testing, Alanine Dipeptide, which still represents a challenge when one biases a very sub-optimal CV. Then, we test one of the standard protein-ligand binding systems, Trypsin-Benzamidine, where the difficulty for a sampling method lies in achieving a subtle balance between being aggressive enough to overcome the many hidden kinetic barriers, and delicate enough not to end up in unwanted conformational states or even unfold the protein. At last, we simulate the canonical protein folding system Chignolin, where an aggressive biasing method is better suited to trigger global folding-unfolding events. Furthermore, we show that our new algorithm is able to provide at no additional cost significant features of the process such as entropy, enthalpy, and the melting temperature of the protein. All examples are compared to analogous PT-MetaD simulations in the Well-Tempered ensemble (PT-WTE-MetaD)^44,45,54^.

Our results compare very favourably with existing state-of-the-art simulations^23,59,60^. The examples provide a scenario for the intended use that we envision for OneOPES: to efficiently exploit the available resources in the study of real-world applications, striking a balance between the human effort needed to design optimal CVs and the computational effort to run long simulations.

## Methods

OneOPES is an explicit replica-exchange technique, whose framework is designed as a progressive stratification of three different external bias potentials (see Fig. 1), which are gradually layered in a sequence of replicas. Here, we present the optimal strategy that we have devised to study the examples that we propose. This approach simply requires the system-dependent optimisation of three key parameters: the leading bias deposition frequency `PACE`, the estimated energy range to be explored `BARRIER` and the maximum temperature to be reached `TEMP_MAX`. We include a total of 8 replicas per system, but the method can be trivially modified and tuned to include a different number of replicas. OneOPES includes two distinct enhanced sampling techniques, i.e. OPES Explore and OPES MultiThermal, and is entirely implemented in the popular open-source plug-in PLUMED2^61^. Below, we give an overview of the main features of each of the techniques and then further discuss their combined use in OneOPES.

**Figure 1.**
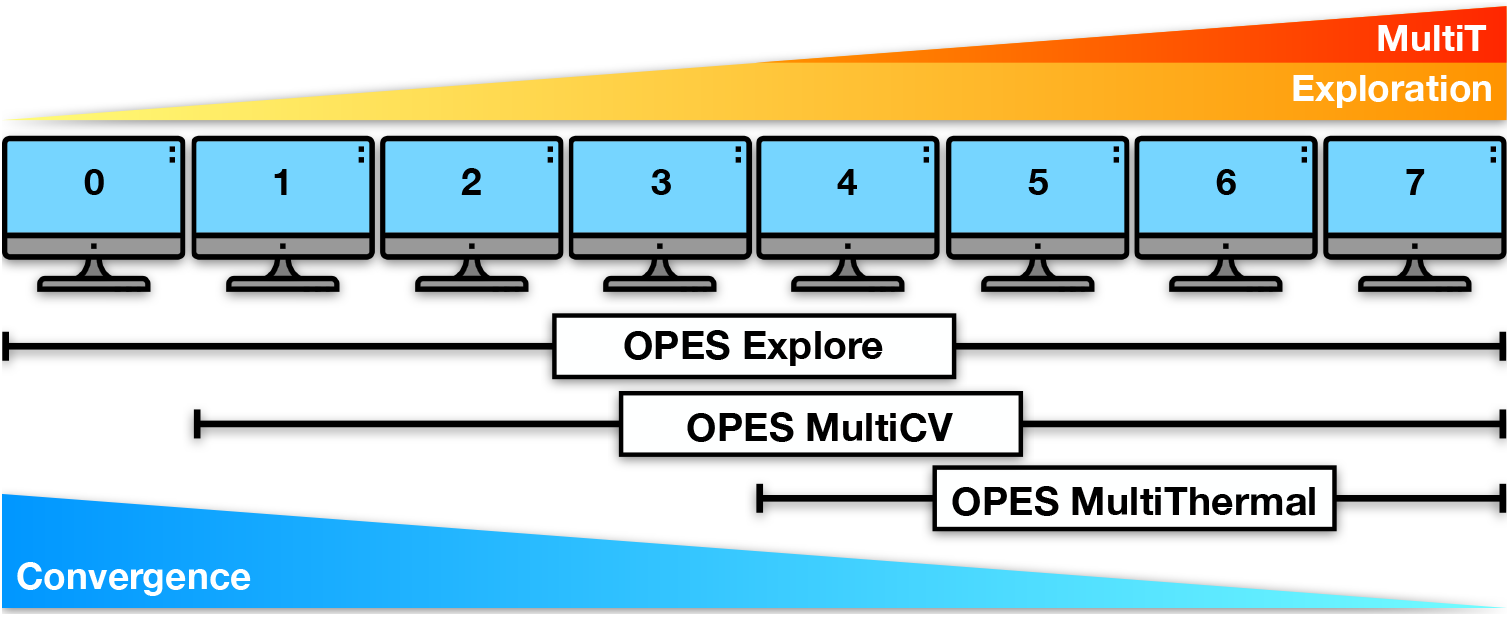
Schematic representation of the OneOPES replica exchange method. Replica **0** only includes one OPES Explore bias potential and is the most convergence-focused replica, while replica **7** is the most exploration-focused one as it may include both extra OPES Explore potentials on additional CVs and OPES MultiThermal with the highest thermal excursion.

**Figure 2.**
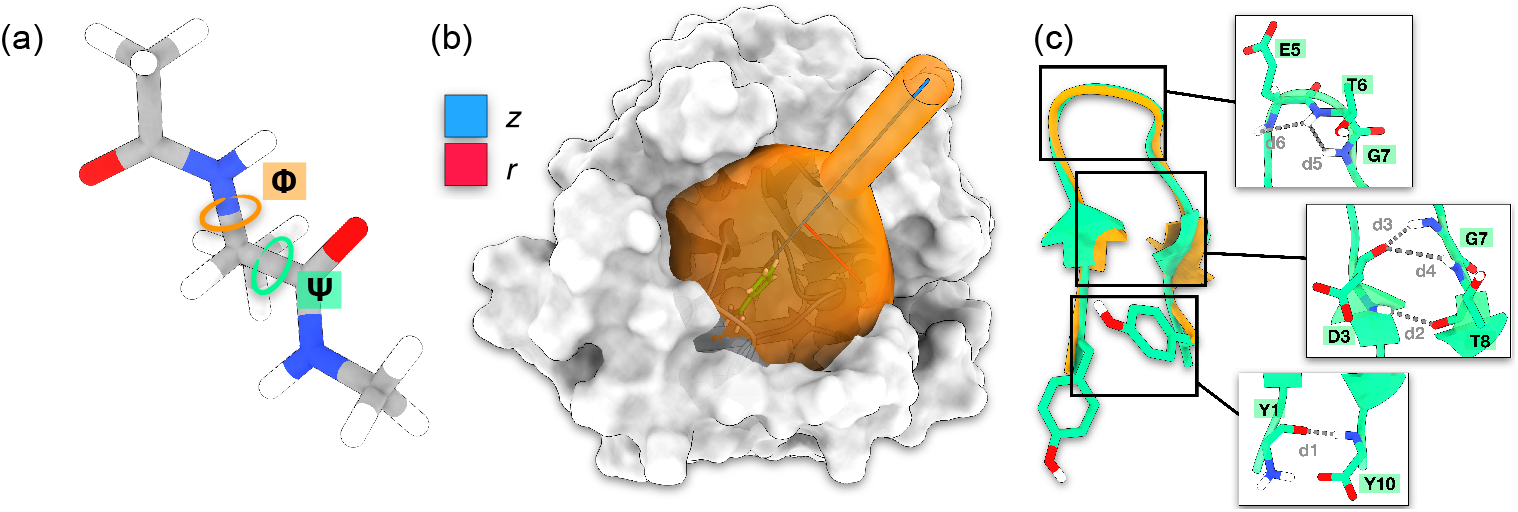
Graphical depiction of the systems that we investigate with OneOPES. In (a), we show Alanine dipeptide with the *ϕ* and *ψ* dihedrals coloured in orange and green, respectively. In (b), we present the Trypsin-Benzamidine complex, with the height *z* and the radius *r* of the funnel that we employ as CVs coloured in blue and red, respectively. In (c), we show the Chignolin miniprotein. We superimpose the Wild-Type structure in orange with the double mutant CLN025 that we simulate in green. The residues and the intraprotein contacts included in the HLDA CV are displayed in the panel insets and highlighted through grey dashed lines.

The first layer of OneOPES is represented by OPES Explore^58^ that is the main simulation bias to drive transitions and reweight trajectories. In each example, all replicas use the same input parameters but, at variance with implementations such as MW^43^, each replica builds its own local bias potential. OPES Explore is a recent evolution of MetaD aimed at making the system sample a broadened target probability distribution 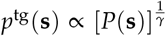 called the Well-tempered distribution, where *P*(**s**) is the unbiased marginal distribution, **s** are the chosen CVs and *γ* is the bias factor that controls the broadening. To achieve this, Gaussian Kernels are used to reconstruct *p*^tg^(**s**) which in turn determines the bias potential *V*(**s**) through a recursive strategy that at step *n* reads

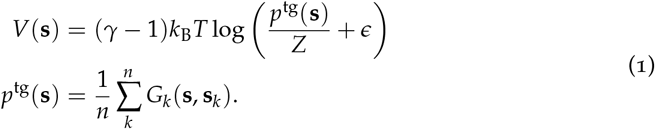

where *k*_B_ is the Boltzmann constant, T the temperature set by the thermostat, *Z* a normalisation factor, and *G*_*k*_ (**s, s**_*k*_) is the Gaussian kernel deposited at step *k*. The initial Gaussian kernel width `SIGMA` is typically the standard deviation of **s** in the initial basin. The bias factor is set by default through *γ* = Δ*E*/(*k*_B_ *T*). The regularisation term *ϵ* is a function of Δ*E* through the relation 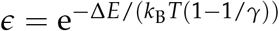

Δ*E* sets a limit on the maximum amount of bias energy that OPES can inject in the system. It should be larger than the expected maximum free energy barrier of the process under investigation so that the bias potential is able to drive transitions away from and back towards the initial basin, while it should not be too large, so that the system does not trigger transitions to high energy states that are irrelevant to the process of interest and may be difficult to reverse. While setting a correct `BARRIER` is important, we have observed that the performance of OneOPES is not too sensitive to this parameter and the choice of a reasonable `BARRIER` value leads to well-converged results in the diverse set of systems studied here.

The frequency at which one deposits Gaussian kernels to update the *p*^tg^(**s**) estimate is another significant parameter called `PACE`. The OPES Explore bias potential is by construction more coarse and changeable in time than the one built in standard OPES^56^. It was shown to guarantee a quick and intensive phase space exploration^26,58,62^. We have found that a sensible choice is to set a `PACE` slower than values typically used in other CV-based enhanced sampling schemes, of the order of thousands of timesteps, and to attempt coordinate exchanges between replicas on a quicker basis, in our case tenfold faster. This way, the system is encouraged to relax in between bias deposition updates and exchange between replicas, gaining access to new conformations through temperature-triggered transitions. Because of this setting, we recommend against using OPES’s default adaptive `SIGMA` scheme that changes in time the Gaussian kernels width according to the CVs’ dynamics. The sudden appearance of different configurations can make in some cases the sigma too large.

The second layer of OneOPES is embedded in replicas **1**-**7** and is represented by auxiliary OPES Explore bias potentials applied on a number of different extra CVs. The role of these OPES MultiCV biases is to promote transitions along transversal DOFs. For these complementary bias potentials, we have found that a low `BARRIER` and a slow `PACE` lead to the best performance. As discussed in Ref. 48, converging a bias potential on individual independent CV is equivalent to converging a fully multidimensional bias, but much faster. To maximise the effectiveness of OPES MultiCV, we recommend choosing a diverse set auxiliary CVs that is largely decoupled from the main CVs. In the examples, we introduce three extra CVs, with the bias on the first one appearing in replicas **1**-**7**, the second one in replicas **2**-**7** and the third one in in replicas **3**-**7**. This progressive introduction of the extra bias potentials is not fundamental but is beneficial to the exchange rate between replicas.

The third layer is represented by OPES MultiThermal^57^ that is aimed at further improving the convergence capabilities of the strategy in the presence of suboptimal CVs. In the examples, it is included in replicas **4**-**7**. By enhancing the fluctuations of the potential energy *U*, OPES MultiThermal allows the system to sample the multicanonical ensemble corresponding to temperatures *T*_*j*_ with *j* = 1, …, *N*_T_ in the range [*T*_min_,*T*_max_]. The free energy difference between each temperature Δ*F*(*T*_*j*_) is iteratively updated while the bias potential is built. At step *n*, the bias potential is as follows

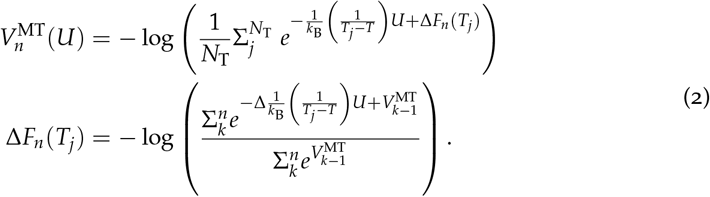

By effectively heating and cooling the system, OPES MultiThermal helps to overcome free energy barriers in a similar fashion to Parallel-Tempering techniques. It is particularly useful to accelerate the sampling along unknown DOFs that are not taken into consideration by the CVs **s**. In the examples, we update the OPES MultiThermal 100 times faster than the main OPES Explore `PACE` so that the OPES MultiThermal goes to convergence faster and grants temperature-triggered transitions. An optimal temperature range must strike a balance between being broad enough to significantly enhance configuration sampling and not too broad to driving the system towards unwanted high energy states.

All in all, at any given step, each replica *i* presents a potential energy *U*_*i*_ (*x*_*i*_) and a total bias potential 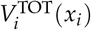 given by the sum of the biases that are applied to it. If we define 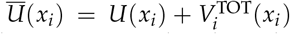 swaps of coordinates between neighbouring replicas *i* and *j* are attempted and regulated by the Metropolis–Hastings algorithm with an acceptance of

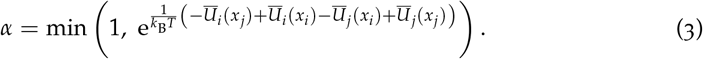

Large temperature intervals [*T*_min_, *T*_max_] applied directly on replica **4** can hamper exchanges with replica **3** and act as an exchange bottleneck. To alleviate this problem, one can use a gradual increase in the temperature range between replicas as we will show in some of the examples. The exchange frequency between replicas must be typically higher than a threshold of about 10% to ensure the diffusion of explorative replicas down to convergence ones and prevent the appearance of exchange bottlenecks. Furthermore, continuous trajectories in which coordinate exchanges are reverted should still display a complete sampling of the phase space. In the Supporting Information (SI) we look into these two aspects and do not see the appearance of exchange bottlenecks in the examples and observe that continuous trajectories sample well the phase space.

The combination of the OneOPES bias potentials facilitates the creation of a double gradient along our replicas framework, i.e. an “exploration gradient” (from **0** to **7**) and a “convergence gradient” (from **7** to **0**), see Fig. 1. The most explorative replicas act as a generator of transitions between different states, that are distilled towards the convergent replica **0**. If relevant slow DOFs are not included in the main CVs, relevant transitions can still occur thanks to the exploratory power of OPES MultiCV or the energy fluctuations of OPES MultiThermal. These transitions allow to visit a system’s free energy minima possibly bypassing the correct transition state region. Therefore, the resulting free energy surface (FES) from these calculations would reproduce well the minima, but would not be able to reliably describe the transition state region. Nevertheless, as we will see in the examples, the better the CV used, the better reconstruction of the FES in all regions, including the transition state.

### Computational details

All calculations are run using the GROMACS 2022.5 engine^63^ patched with the PLUMED 2.9 plugin^61^ with the exchange algorithm implemented in Ref.^47^. Further simulation details and the simulations’ computational cost are provided in the SI. We study three standard biophysical examples: conformational changes in Alanine dipeptide, protein-ligand binding in Trypsin-Benzamidine, and protein folding in the Chignolin miniprotein. To highlight the impact of using extra CVs we perform both simulations where we bias them and simulations where we do not include them. We will call the former strategy *OneOPES MultiCV* and the latter one *OneOPES*. Furthermore, to compare our strategy with one of the standard enhanced sampling techniques in the field, we include PT-WTE-MetaD simulations tuned to replicate as closely as possible the OneOPES ones. In each case, we perform 5 independent simulations and calculate the corresponding free energy difference as a function of simulation time through reweighting, by using as a weight in OneOPES the OPES Explore instantaneous bias or the value of the bias normalised by the reweighting factor^64^ *c*(*t*) in PT-WTE-MetaD. In the reweighting procedure, we use the most convergence-focused replica **0**, from which we discard the initial 10% of the trajectory.

We compare the average and standard deviation of the independent simulations free energy difference and FES with highly converged results. In Alanine dipeptide the reference free energy difference 8.9 *±*0.1 kJ/mol is calculated from a 100 ns OPES simulation where both the *ϕ* and *ψ* dihedral angles are biased and a 10-block analysis is performed.

In Trypsin-Benzamidine the reference Δ*F* = 26.6*±*0.3 kJ/mol is taken from the extensive calculations presented in Ref. 59. In Chignolin the reference Δ*F* = 3.6*±*0.4 kJ/mol is calculated with a 10-block analysis from the 100 *μ*s trajectory from Ref. 65.

In Alanine Dipeptide, the main CV that we bias is the suboptimal *ψ* angle, in Trypsin-Benzamidine we bias the funnel^66^ axis *z* and radius *r*, in Chignolin we bias a harmonic linear discriminant analysis (HLDA) CV based on six interatomic contacts within the protein^18,67^. The extra CVs that we choose to bias are: three distances between heavy atoms in Alanine dipeptide; three water coordination sites in Trypsin-Benzamidine; a water coordination site, the gyration radius and a contact between the termini in Chignolin. Additional details about the extra CVs are provided in the SI.

The OPES Explore `BARRIER` parameter is 50 kJ/mol in Alanine dipeptide and Chignolin, while it is 30 kJ/mol in Trypsin-Benzamidine. The deposition `PACE` is 5000 simulation steps in Alanine dipeptide, 10000 steps in Trypsin-Benzamidine and 100000 steps in Chignolin. The OPES Explore `PACE` determines other parameters such as the OPES MultiThermal `PACE` that is set to be 100 times faster, the replica exchange frequency that is 10 times faster and, when present, the OPES Explore MultiCV `PACE` that is 2 times slower. The `BARRIER` parameter on the MultiCV biases is always 3 kJ/mol. In OPES MultiThermal, the `TEMP_MAX` reached in replicas **4**-**7** in Alanine dipeptide is 600K, in Trypsin-Benzamidine it is respectively [310 K, 330 K, 350 K, 370 K] and in Chignolin [350K, 365 K, 380 K, 400 K].

In PT-WTE-MetaD, as customary, we initially perform short simulations to bring the WTE bias to equilibrium (see SI). Then, we add a MetaD bias potential to all replicas on the same CVs as OneOPES and slow down the bias deposition of the WTE MetaD. In all simulations, a WT-MetaD is performed on the same sub-optimal CVs as above with a Gaussian Kernels `HEIGHT` of 1.5 kJ/mol, a `PACE` of 500 steps and a replica exchange frequency of 10000 steps. In Alanine dipeptide the `BIASFACTOR` is 20 and the thermostat in the explorative replicas is set to [357 K, 425 K, 506 K, 600 K]. In Trypsin-Benzamidine the `BIASFACTOR` is 15 and the thermostat in the explorative replicas is set to [305 K, 319 K, 334 K, 350 K]. In Chignolin the `BIASFACTOR` is 20 and the thermostat in the explorative replicas is set to [352 K, 364 K, 377 K, 390 K].

## Results

### Alanine Dipeptide

Alanine dipeptide in vacuum is a prototypical system, routinely used in the early phase of method development^1^. The system presents two conformational states that depend upon dihedral angles *ϕ* and *ψ*. In this regard, *ϕ* is a nearly-ideal CV to capture the conformational change as it distinguishes well the two states and the corresponding transition state. Conversely, *ψ* is far from ideal as it barely distinguishes the states and it is almost orthogonal to the transition state.

At first, we perform calculations biasing *ϕ* to verify the strategies’ convergence in combination with optimal CVs. All simulations converge to the exact result within a few nanoseconds (see Fig. S1). Moreover, we notice that the FES is well described in all regions, including the transition state. This is not surprising and confirms that using optimal CVs in enhanced sampling simulations, OneOPES included, is the best route to obtain high quality well-converged results in short simulation times.

Regrettably, optimal CVs are hard to come by in realistic systems. To replicate the effect of using bad-quality CVs, we perform a more demanding test on Alanine Dipeptide by biasing the suboptimal CV *ψ*. In a recent paper^23^, it was shown that a 5 *μ*s enhanced sampling simulation where the authors biased *ψ* is capable of triggering just a handful of transitions and does not reach convergence. In the same paper, a 50 ns OPES MultiThermal simulation represents an improvement, as it produces a converged but fairly noisy Δ*F* from its still limited number of transitions.

In Fig. 3 we show the free energy difference and the FES from five independent simulations performed with different replica exchange methods. In panels (a) and (d) we show the PT-WTE-MetaD results where, after an initial phase in which the average Δ*F* between simulations displays a rather large standard deviation, for a longer simulation time it tends to roughly agree with the expected value, albeit being slightly down-shifted. In panels (b) and (e), we use OneOPES and observe an improved match between mean values and exact results, with a variance between independent simulations that shrinks in time and a mean Δ*F* that gets closer to the ideal one.

**Figure 3.**
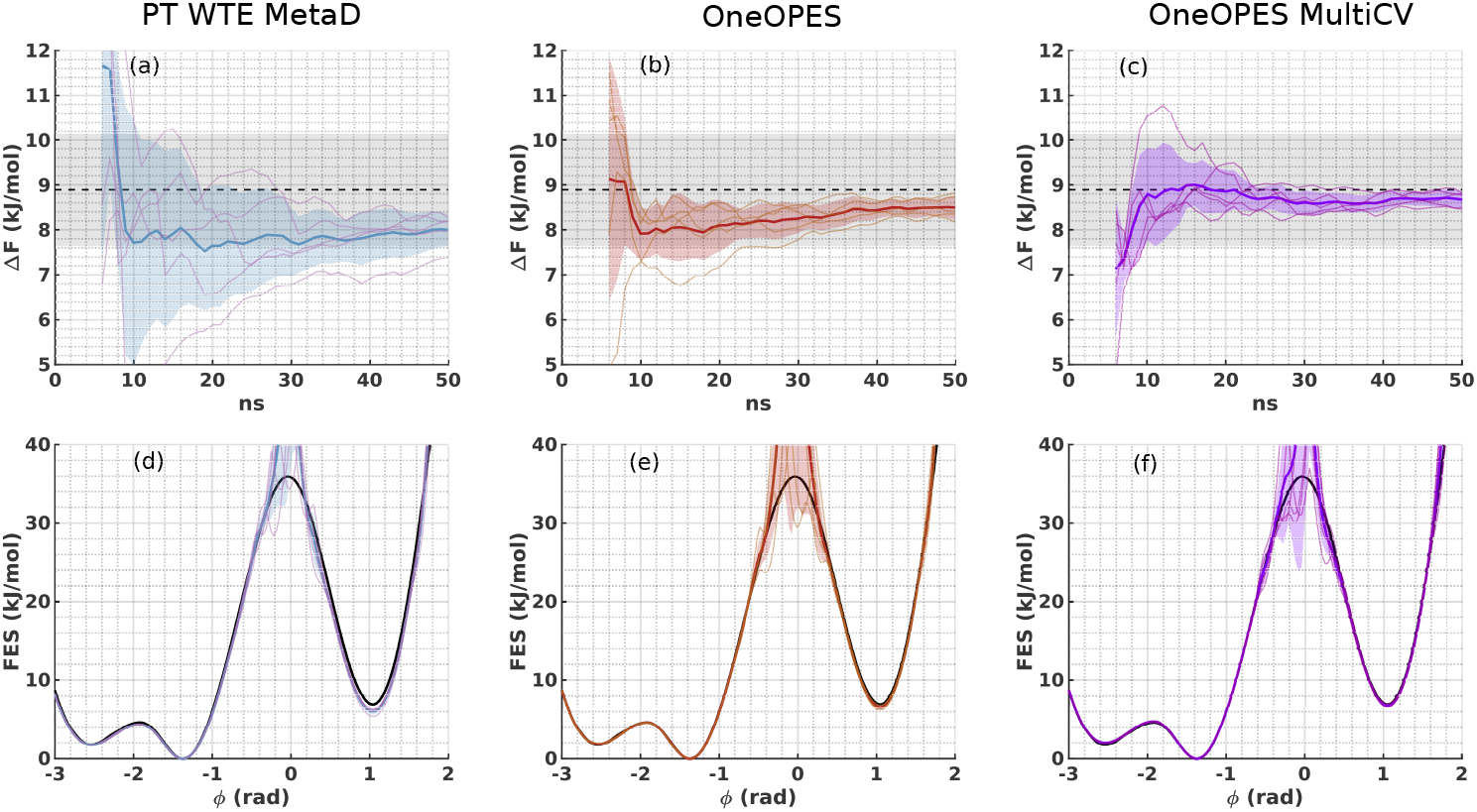
Alanine Dipeptide set of 5 independent simulations where we bias the sub-optimal CV *ψ* with PT-WTE-MetaD (panels (a) and (d)), OneOPES (panels (b) and (e)) and OneOPES MultiCV (panels (c) and (f)). In (a-c), we show the average Δ*F* in time with a dark blue, dark red and dark purple solid line and their standard deviation in semitransparent regions in light blue, light red and light purple, respectively. Δ*F* values corresponding to individual simulations are shown with solid thinner lines. The expected Δ*F* is indicated by a dashed black line with an tolerance error of 0.5 *k*_B_ *T* in shaded grey. In (d-f), we show the one-dimensional FES reweighted over *ϕ* after 50 ns. The same colour scheme applies as in panels (a-c).

Finally, in panels (c) and (f) we present the results of OneOPES MultiCV. While the convergence in the long term is similar to that of OneOPES, in the short term (*≈* 10 ns) the presence of the additional bias from the extra CVs helps all the independent simulations to reach a mean Δ*F* value closer to the expected one and that converged value is kept until the end of the simulation. In this example, the OneOPES scheme is able to drive the system back and forth between states and obtain a converged FES even when coupled with ineffective CVs. Example trajectories are provided in Figs. S4-S6 in the SI. We wish to point out that, at variance with PT-WTE-MetaD, the instantaneous value of the bias in OneOPES tends to reach a correct quasi-static value earlier, which in turn guarantees a faster and more robust convergence.

In all simulations we notice that, while the free energy in the main basins closely matches the exact ones, the same is not true for the barriers associated to the transition regions. Like in Parallel Tempering schemes, this occurs because the transition regions are less effectively sampled and are often skipped over through the exchanges with the the explorative replicas. In Figs. S4-S6 in the SI, we compare the 2-dimensional FES of replica **0** and **7** and, as expected, we notice that, when biasing the suboptimal CV *ψ*, the most explorative replica **7** better samples the transition state region.

### Trypsin-Benzamidine

A more arduous test is the ligand-binding benchmark Trypsin-Benzamidine. While this system has been routinely used for years as a benchmark for ligand-binding methods^66,68–80^, it is far from trivial and still offers a significant challenge. High-resolution crystallographic experiments have recently demonstrated that individual water molecules play a crucial role in the system’s binding/unbinding process^81^, so the introduction of specialised water-focused CVs proved decisive in bringing simulations to convergence in our recent work^59^. While the information provided by such CVs is invaluable, it is nevertheless an unfeasible task to replicate its development and optimisation in a high-throughput context.

In this paper, we pursue a different approach and we simulate the system in combination with standard CVs that only capture the motion of the ligand with respect to the binding pocket and neglect the water dynamics (see SI). These CVs are clearly not optimal for the problem at hand but represent a more suitable option for future applications where one wants to extract binding free energies of sufficient quality from a number of systems, without focusing on a case-by-case CV optimisation.

The free energy difference and the FES from PT-WTE-MetaD (panels (a) and (d)) show a marked shift compared to the expected result from Ref. 59. In SI Fig. S8(b) we show the normalised bias dynamics in an example trajectory and notice that it displays large fluctuations until it stabilises itself towards the end of the simulation. To investigate if discarding more of the initial portion of the data would improve the binding free energy estimate, we repeat the reweighting procedure by discarding respectively 20%, 40% and 60% of the trajectory and show the results in Fig. S11. It is apparent that the closest agreement with the expected free energy is reached by discarding at least 60% of the data, indicating that PT-WTE-MetaD simulations would eventually provide accurate free energy estimates for this system, but, to achieve so, they would require a rather long sampling time.

In Fig. 4(b,d) we show the corresponding results of OneOPES and observe a notable improvement in both the agreement with the expected result and the speed at which it is achieved. Fig. S9(b) reveals that the bias here reaches a quasi-static condition earlier in the simulation with respect to PT-WTE-MetaD. Furthermore, the gentle phase space exploration granted by OneOPES is crucial in systems such as Trypsin-Benzamidine, where, on the other hand, an aggressive bias may lead to irreversible local conformational changes and ultimately produce incorrect free energy profiles.

**Figure 4.**
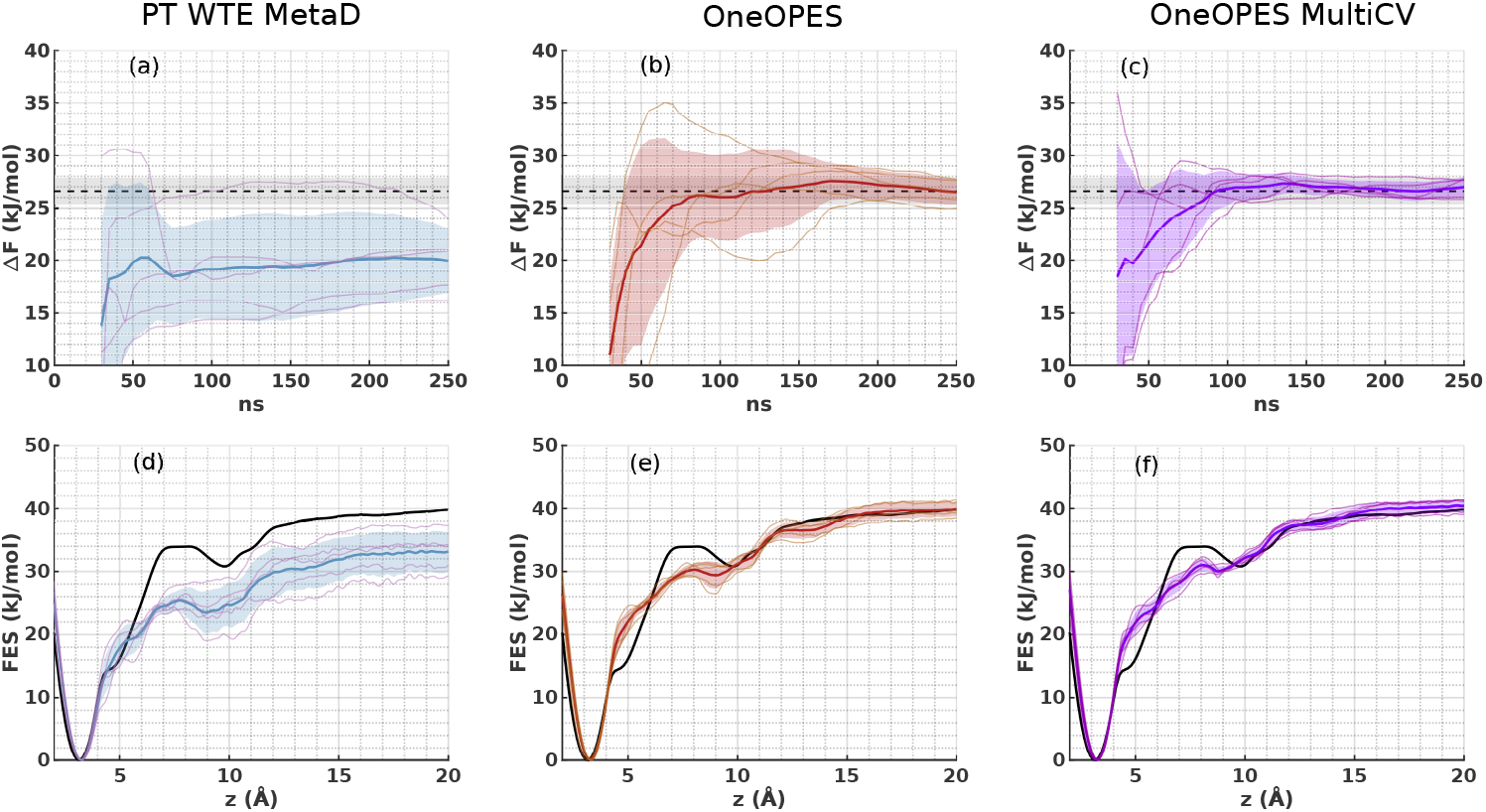
Set of 5 independent Trypsin-Benzamidine simulations where we bias the funnel coordinates *z* and *r* ^66^ with PT-WTE-MetaD (panels (a) and (d)), OneOPES (panels (b) and (e)) and OneOPES MultiCV (panels (c) and (f)). In (a-c), we show the average Δ*F* in time with a dark blue, dark red and dark purple solid line and their standard deviation in semitransparent regions in light blue, light red and light purple, respectively. Δ*F* values corresponding to individual simulations are shown with solid thinner lines. The expected Δ*F* is taken from Ref.^59^ and is indicated by a dashed black line with an tolerance error of 0.5 *k*_B_ *T* in shaded grey. In (d-f), we show the one-dimensional FES reweighted over *z* after 250 ns. The same colour scheme applies as in panels (a-c).

A critical DOF in the Trypsin-Benzamidine binding process is represented by the long-lived water molecules directly affecting the binding free energy. In OneOPES such water molecules are accelerated by OPES MultiThermal. In OneOPES MultiCV we choose to explicitly bias the water coordination around three relevant hydration spots among the ones pointed out in Ref. 59, i.e., one around the ligand, one around the binding site, and one at the entrance of the water reservoir cavity (see SI). The resulting free energies presented in Fig. 4(c,e) are even more accurate than the ones from OneOPES and, remarkably, all the simulations independently reach a converged and stable Δ*F* after about 100 ns. Corresponding trajectories are shown in the SI and in Fig. S10.

### Chignolin

Protein folding are complex examples to study with CV-based enhanced sampling methods due to the difficulty for CVs to capture processes characterised by a sequence of intermediate metastable states^82^. A further source of complications is the fact that a protein’s folded and unfolded states are intrinsically different, as the former is enthalpy dominated while the latter is entropy dominated. Developing CVs that are optimal in capturing the whole folding/unfolding transition is a very complex task that today can be accomplished only in the simplest cases^23^.

We focus on such a relatively simple case and use OneOPES to study the folding of CLN025, a double mutant (G1Y, G10Y) of the Chignolin miniprotein^83,84^, a system largely employed in the last decade as a benchmark to study fast-folding proteins through both long unbiased molecular dynamics simulations^65^ and more recently enhanced sampling simulations^23,29,50,57,67,85–87^. We will employ a rather simple CV that is based on the linear combination of six interprotein contacts whose weights are obtained through harmonic linear discriminant analysis (HLDA)^18,67^ (see SI for additional details). Following the scheme used in the previous examples, we perform 5 PT-WTE-MetaD, OneOPES and OneOPES MultiCV simulations, and compare their resulting free energies with the reference one from Ref.^65^.

As visible in Fig. 5 (a,d), PT-WTE-MetaD simulations converge within 0.5 *k*_B_*T* from the expected result in about 150 ns, but the estimated error does not shrink for a longer simulation time and the mean Δ*F* tends to marginally drift away from the expected value. The corresponding OneOPES results in Fig. 5 (b,e) show analogous behaviour.

**Figure 5.**
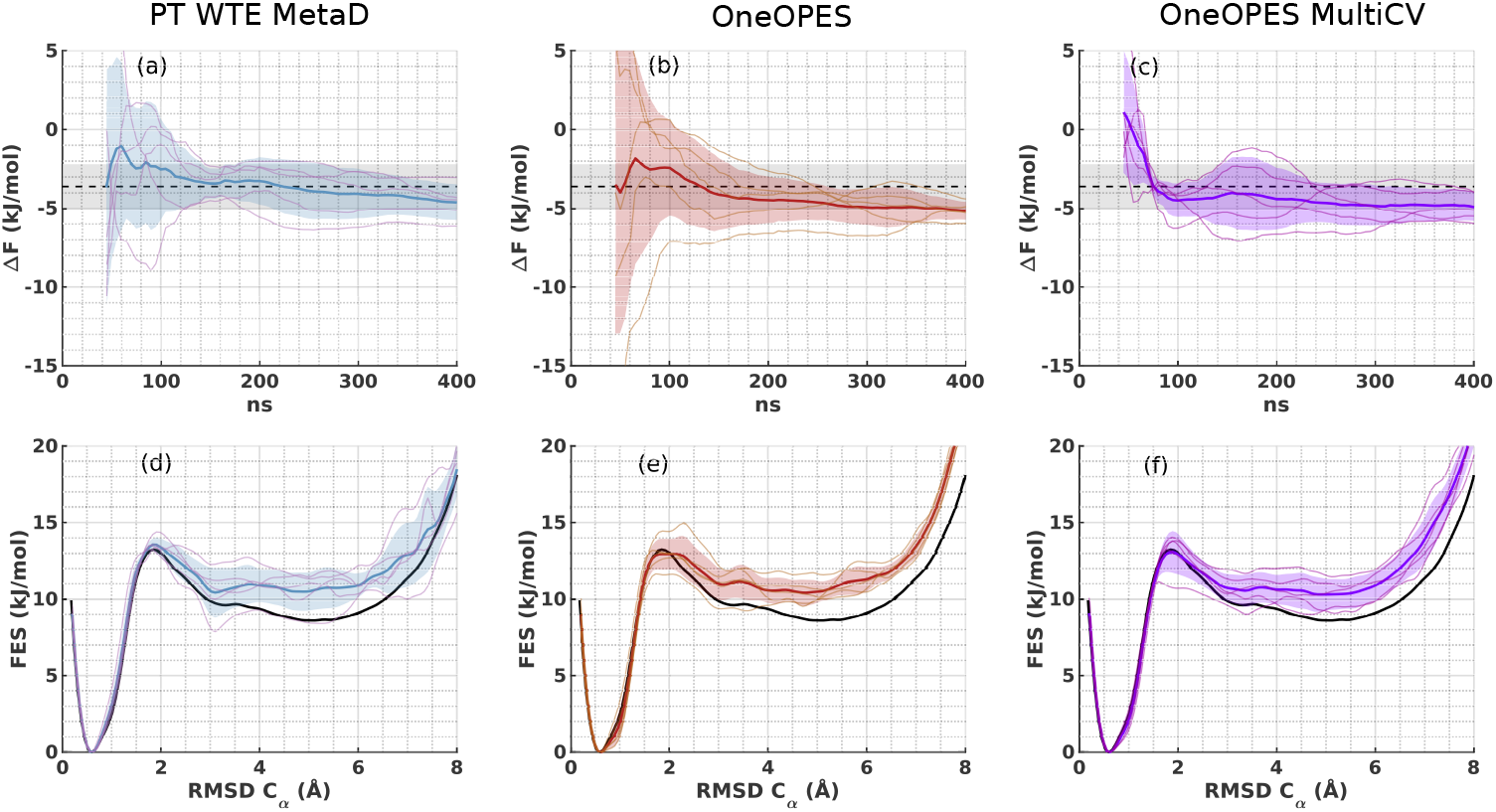
Set of 5 independent Chignolin simulations where we bias the HLDA CV^18,67^ with PT-WTE-MetaD (panels (a) and (d)), OneOPES (panels (b) and (e)) and OneOPES MultiCV (panels (c) and (f)). In (a-c), we show the average Δ*F* in time with a dark blue, dark red and dark purple solid line and their standard deviation in semitransparent regions in light blue, light red and light purple, respectively. Δ*F* values corresponding to individual simulations are shown with solid thinner lines. The expected Δ*F* is taken from Ref.^65^ and is indicated by a dashed black line with an tolerance error of 0.5 *k*_B_ *T* in shaded grey. In (d-f), we show the one-dimensional FES reweighted over the RMSD C*α* after 400 ns. The same colour scheme applies as in panels (a-c).

In OneOPES MultiCV we add extra bias potentials on a diverse set of CVs, i.e. a water coordination site, the protein gyration radius and the contact between the termini. In Fig. 5 (c,e), we see that the mean Δ*F* converges faster than PT-WTE-MetaD and OneOPES, but it is still slightly shifted. In Fig. S12-S14 in the SI, we show typical trajectories and we point out that the bias dynamics in all the simulations hardly reaches a quasi-static condition and displays especially large fluctuations in the PT-WTE-MetaD case (Fig. S12(b)).

We then decide to investigate two further scenarios. In one, we replace the HLDA CV in OneOPES with two well-known highly suboptimal CVs for driving protein folding, i.e. the RMSD on the C*α* and the radius of gyration. The resulting free energy results shown in Fig. S15(a,d) in the SI display a much noisier and less converged behaviour than those performed on the HLDA CV. The mean Δ*F* between simulations has a rather large standard deviation but it still is in qualitative agreement. Corresponding trajectories in Fig. S16 displays less unfolded to folded events, in line with what is expected from a less efficient CV.

The second scenario that we investigate is one where we increase the aggressivity of OneOPES by doubling the `BARRIER` parameter in OPES Explore (see SI). The resulting Δ*F* in Fig. S15(b,e) in the SI is in good agreement with the expected result and does not display a shift anymore. As expected, this setting makes the bias dynamics even more noisy, as shown in Fig. S17(b). In this more aggressive case, the use of extra CVs in OPES MultiCV does not seem to bring any benefit (Fig. S15(c,f)). We believe that this perhaps counter intuitive behaviour is largely due to the shortcomings of the HLDA CV to comprehensively capture the complexity of protein folding and the safest route to follow would be to craft an improved CV that includes more information about the process.

One of the advantages of using OPES MultiThermal is possibility to evaluate physical properties in a range of temperatures away from the temperature set by the thermostat through a reweighting procedure^57^. Therefore, we exploit this feature and use replica **6** from the OneOPES MultiCV simulations on the HLDA CV to estimate the system’s folding Δ*F* between 340K and 360K (see Fig. S19 in the SI). Through this procedure, we estimate the melting temperature of Chignolin to be 405*K ±* 9*K*, which is in reasonable proximity with the value of 381*K* with a 68% confidence interval of 361 *−* 393*K* from Ref. 65.Moreover, by performing a linear fit of the Van ‘t Hoff equation Δ*F* = Δ*H − T*Δ*S* we can also estimate the enthalpy Δ*H* and entropy *−T*Δ*S* of folding, which are *−*32.2 *±*3.1 kJ/mol and 27.0 *±*2.5 kJ/mol respectively (see Tab. S4 for more information).

## Conclusions

Collective variable-based enhanced sampling MD simulations rely on optimal CVs that approximate the reaction coordinate and encapsulate all the relevant slow DOFs so that they can accelerate their sampling. A serious drawback of these approaches is the effort required to define optimal CVs that capture complex processes such as folding, the flexibility of a receptor^88,89^ or the role of water at a ligand/protein’s interface^90–100^. Methods based on machine learning are increasingly successful in providing optimal CVs, but they need significant amount of data that is not always available^10,12,14–29^.

As a result, the use of suboptimal CVs is often unavoidable. To address this problem, we propose a novel replica-exchange framework named OneOPES, based on the combination of the recently developed OPES Explore and OPES MultiThermal methods. OneOPES is able to compensate some of the CVs’ shortcomings by setting up a hierarchy of replicas in convergence and explorative power, making out-of-equilibrium barrier crossings occur mostly on explorative replicas and letting configurations exchange towards convergence-focused replicas. We have shown that OneOPES can consistently recover the free energy of the increasingly complex examples that we benchmarked (i.e. Alanine dipeptide, Trypsin-Benzamidine, and Chignolin) within an error of 0.5 *k*_B_ *T* at a reasonable computational cost, even in combination with suboptimal CVs. At the same time, it unlocks the possibility to infer thermodynamical properties of the system under investigation such as the enthalpy, the entropy, and the melting temperature.

We emphasise that although OneOPES is very effective and represents a significant advance over other OPES and Metadynamics variants, we do not expect it to be miraculous in combination with very poor quality CVs. If the CVs are not relevant or inadequate for the problem at hand, OneOPES would still fail to converge. Still, in combination with reasonable CVs that distinguish the important states of the system but are not fine-tuned (since crucial but hard-to-capture DOFs are missing from the chosen set of CVs) the correct free energy landscape can be still recovered with a reasonable amount of sampling. Furthermore, the inclusion of a handful of extra CVs in additional perturbative biases shows a further promising route for improvement in the speed at which converge is reached.

We expect that the OneOPES approach will fit especially well the study of complex biophysical phenomena, ranging from conformational changes to ligand binding. In particular, we believe that our results endorse OneOPES as a valuable tool that can be safely used beyond benchmark systems in the study of complex and so far unexplored systems whose optimal CVs are still unknown. In venturing in this direction, OneOPES can be easily combined with state-of-the-art machine-learning CV design techniques, stretching to the limit the boundary of the systems that can be studied by modern computational techniques.

## Data Availability

OneOPES is implemented in PLUMED^61^, from version 2.8 onward, in combination with GROMACS^47,63,101–105^. The input files to replicate all the simulations and the corresponding analysis scripts can be found on the PLUMED NEST repository^106^ https://www.plumed-nest.org/eggs/23/011/.

## Supplementary Information Available

## Acknowledgements

We acknowledge PRACE and the Swiss National Supercomputing Centre (CSCS) for large supercomputer time allocations on Piz Daint, project IDs: pr126, s1107, s1169, s1228. We acknowledge the Swiss National Science Foundation and Bridge for financial support (projects number: 200021_204795 and 40B2-0_203628). We thank Michele Parrinello, Umberto Raucci, Dhiman Ray, Michele Invernizzi, and Ioannis Galdadas for helpful discussions.

