## Supplementary Information for "OneOPES, a combined enhanced sampling method to rule them all"

#### Supporting Computational Details

To calculate free energy surfaces and differences in OneOPES simulations, we apply the standard reweight procedure from Ref. [1] to replica **0**, the most convergence-focused one. As customary, we discard the initial portion of the trajectories where the bias potential may still be out-of-equilibrium. Here this portion amounts to 10% of the total simulation time.

##### Alanine Dipeptide

The 3D structure of Alanine Dipeptide was built from scratch, capping the N-terminus and C-terminus with an acetyl and a methyl-amino protecting group, respectively. The Amber ff99sb-ildn force field was employed for the MD simulation [2], which was run using the GROMACS 2022.5 engine [3] patched with the PLUMED 2.9 plugin [4]. The MD simulation was run in vacuum with a timestep of 2 fs. The temperature was kept in check through the V-rescale thermostat [5]. Periodic boundary conditions were applied, and the particle-mesh-Ewald (PME) method was used to treat long-range electrostatic interaction [6]. For short-range interactions, a cut-off distance of 1.0 nm was applied. Two leading CVs have been employed on OPES Explore to estimate the free-energy, i.e. either the torsional angle $\phi$ or the torsional angle $\psi$ (see Fig.2a). On OPES MultiCV, we employed the following auxiliary CVs:

- Acetyl's C$\beta$-Methylamine's C$\alpha$ (**d1**)
- Acetyl's Oxygen-Methylamine's Nitrogen (**d2**)
- Alanine's Nitrogen-Alanine's Oxygen (**d3**)

Regarding the replicas scheme, we used 8 replicas in parallel according to Fig. 1, whose frequency of exchange was 500 integration steps. All replicas contained an OPES Explore bias with a $\Delta E$ of 50 kJ/mol, a SIGMA of 0.2 and a PACE of 5000 steps. Additionally, replicas **1-7** underwent OPES Explore with a $\Delta E$ of 3.0 kJ/mol on the above-mentioned auxiliary CVs. Each CV was embedded into a separate layer of OPES Explore, i.e. 1 layer for replica **1** (sampling **d1** with a SIGMA of 0.02 nm), 2 layers for replica **2** (sampling **d1** and **d2** with a SIGMA of 0.015 nm), and 3 layers for replicas **3-7** (sampling **d1**,**d2**, and **d3** with a SIGMA of 0.01 nm). All MultiThermal replicas explore the temperature range [300K, 600K]. The total simulation time per replica is 50 ns.

The PT-WTE-MetaD simulations were preceded by a preliminary part where the WTE bias was brought to convergence. This simulation lasted 2 ns and had replicas **0-3** unbiased, while replicas **4-7** included a MetaD bias on the system internal energy with PACE 1000 steps, SIGMA 10 kJ/mol, HEIGHT 0.2 kJ/mol and bias factor 10. The thermostat in such replicas was set to [357 K, 425 K, 506 K, 600 K]. After this preliminary simulation, the MetaD bias on the system internal energy was restarted with a PACE of 500000 steps, while a MetaD bias on $\psi$ was introduced in all replicas. Such bias had PACE 500 steps, SIGMA 0.2 kJ/mol, HEIGHT 1.5 kJ/mol and bias factor 20. The replica exchange frequency was every 10000 steps.

In the main text, the results are compared to an ideal 100 ns simulation where we use OPES to bias both $\phi$ and $\psi$ with $\Delta E$ of 60 kJ/mol and a PACE of 500 steps.

#### Trypsin-Benzamidine

The initial coordinates of the Trypsin-Benzamidine complex were retrieved from PDB ID: 2OXS [7]. The ligand-protein complex was inserted into a box filled with 14704 TIP3P water molecules and filled with 7 Chlorine ions to ensure system neutrality. The simulation box is cubic with a side of 77.987 Å.

The Amber ff99sb-ildn force field was employed for Trypsin [2], whereas the ligand's topology was generated with GAFF and converted into a GROMACS-readable format in order to use it in the MD engine GROMACS 2022.5 [8]. The benzamidine topology underwent a quantum-mechanical re-parametrization of the torsional angle between the amidine and the benzene ring, i.e. its energy profile was optimised so that it closely matches the one obtained through a dihedral scan carried out with quantum-mechanical calculations. For more details about the procedure, we refer the interested reader to the paper 9.

Temperature control was applied through the V-rescale thermostat [5], and the pressure was fixed at a reference value of 1 bar thanks to the Parrinello-Rahman barostat [10]. Periodic boundary conditions, short- and long-range interactions were treated as in the *Alanine Dipeptide* MD simulation. To measure the binding affinity of Benzamidine toward Trypsin, we employed a funnel-shaped potential [11], in accordance with the foremost literature in the field [12] (see Fig.2b). The funnel position was set up to cover Trypsin's cavity (i.e. the ligand's binding site), with the cylindrical part pointing towards the bulk solution. The vertical axis of the funnel was also exploited to set up two CVs, i.e. the height of the ligand $z$ and the orthogonal distance (i.e. the radius $r$ of the funnel). Similarly to Alanine Dipeptide, auxiliary CVs have been introduced to increase the quality of the sampling:

- water coordination site (**V9**)
- water coordination site (**H**)
- water coordination site (**G**)

For additional information about the water coordination sites, please see Ref.[13].

To calculate the absolute free energy of binding and factor out the funnel's entropic restraint, one needs to apply a correction to the apparent free energy $F(z)$ that comes from the simulation:

$$\Delta F = -\frac{1}{\beta}\log\left(C^0 \pi R_{\text{cyl}}^2 \int_{\text{B}} \text{d}z \, \exp\left(-\frac{1}{k_{\text{B}}T}(F(z) - F_{\text{U}})\right)\right) \tag{S1}$$

where $C^0 = 1/1660$ Å$^{-3}$ is the standard concentration, $z$ is the distance between the centre of mass of the ligand and the binding site along the funnel's axis, $F(z)$ is the free energy along the funnel axis and $F_{\text{U}}$ is its reference value in the unbound state.

Regarding the replicas scheme, we run 8 replicas in parallel as in the *Alanine Dipeptide* MD simulation, where the replicas underwent OPES Explore with a $\Delta E$ of 30 kJ/mol, a SIGMA of (0.02,0.02) nm and a

PACE of 10000 steps. For the MultiThermal replicas, we selected the following temperature range, i.e. [300K, 310K] for replica **4**, [300K, 330K] for replica **5**, [300K, 350K] for replica **6**, [300K, 370K] for replica **7**. Regarding the auxiliary CVs, each has been embedded into different layers of OPES Explore ($\Delta E$= 3 kJ/mol, PACE= 20000 steps), and progressively distributed along the replicas, i.e. 1 layer for replica **1** (sampling **V9** with a SIGMA of 0.05), 2 layers for replica **2** (sampling **V9** and **H** with a SIGMA of 0.1), and 3 layers for replicas **3-7** (sampling **V9**, **H**, and **G** with a SIGMA of 0.1). The total simulation time per replica is 250 ns.

The PT-WTE-MetaD simulations were preceded by a preliminary part where the WTE bias was brought to convergence. This simulation lasted 20 ns and had replicas **0-3** unbiased, while replicas **4-7** included a MetaD bias on the system internal energy with PACE 500 steps, SIGMA 1000 kJ/mol, HEIGHT 5 kJ/mol and bias factor 50. The thermostat in such replicas was set to [305 K, 319 K, 334 K, 350 K]. After this preliminary simulation, the MetaD bias on the system internal energy was restarted with a PACE of 500000 steps, while a MetaD bias on $z$ and $r$ was introduced in all replicas. Such bias had PACE 500 steps, SIGMA (0.2,0.2) nm, HEIGHT 1.5 kJ/mol and bias factor 15. The replica exchange frequency was every 10000 steps.

In the main text, the results are compared to the state-of-the-art simulation from Ref. [13].

#### Chignolin

The Chignolin miniprotein employed in the present manuscript is a double mutant (G1Y, G10Y) of the Wild Type Chignolin (PDB ID: 1UAO), named *CLN025* [14, 15]. As largely employed in recent literature, the protein was parameterised with the CHARMM22* force field [16], then embedded into a box filled with 1907 TIP3P water molecules and 2 sodium ions to ensure system neutrality. The simulation box is cubic with a side of 39.61 Å.

The MD simulation was run with the MD engine GROMACS 2022.5 [3] with a timestep of 2 fs in the NVT ensemble, where the temperature was kept in check through the V-rescale thermostat at 340 K [5]. Periodic boundary conditions, short and long-range interactions were treated as in the *Alanine Dipeptide* MD simulation. To investigate the folding mechanism of Chignolin, we used a tailored CV, i.e the *Harmonic Linear Discriminant Analysis* (HLDA) CV based on six interatomic contacts within the protein [17, 18] (see Fig. 2(c)). To properly compare our estimate of the FES with data reported in the literature, we reweighted the deposited bias through the *RMSD* CV, computed on the C$\alpha$ atoms of CLN025's residues. For the sake of clarity, the following auxiliary CVs have been envisioned:

- water coordination site located on the average Chignolin's C$\alpha$s position (**bubble**)
- coordination between the N-terminus and C-terminus (**cNC**)
- radius of gyration (**rg**)

Regarding OneOPES, we run 8 replicas in parallel as in the *Alanine Dipeptide* MD simulation, where the replicas underwent OPES with a $\Delta E$ of 50 kJ/mol, a SIGMA of 0.04 and a PACE of 100000 steps. For the MultiThermal replicas, we applied a ramp to the temperature ranges in order to boost the replica exchange acceptance ratio, i.e. [340K, 350K] for replica **4**, [340K, 365K] for replica **5**, [340K, 380K] for replica **6**, [340K, 400K] for replica **7**. Regarding the auxiliary CVs, each has been embedded into different layers of OPES Explore ($\Delta E$= 3 kJ/mol, PACE= 200000 steps), and progressively distributed along the replicas, i.e. 1 layer for replica **1** (sampling **bubble** with SIGMA 0.2), 2 layers for replica **2** (sampling **bubble** and **cNC** with SIGMA of 0.06), and 3 layers for replicas **3-7** (sampling **bubble**, **cNC**, and **rg** with SIGMA of 0.006 nm). The total simulation time per replica is 400 ns.

The PT-WTE-MetaD simulations were preceded by a preliminary part where the WTE bias was brought to convergence. This simulation lasted 10 ns and had replicas **0-3** unbiased, while replicas **4-7** included a MetaD bias on the system internal energy with PACE 500 steps, SIGMA 300 kJ/mol, HEIGHT 5 kJ/mol and bias factor 20. The thermostat in such replicas was set to [352 K, 364 K, 377 K, 390 K]. After this preliminary simulation, the MetaD bias on the system internal energy was restarted with a PACE of 500000 steps, while a MetaD bias on the HLDA CV was introduced in all replicas. Such bias had PACE 500 steps, SIGMA 0.03, HEIGHT 1.5 kJ/mol and bias factor 20. The replica exchange frequency was every 10000 steps.

The simulation presented in Fig. S15(a), is identical to the OneOPES one in the main text, except for the main CVs biased by OPES explore that are the *RMSD* on the C$\alpha$ and the radius of gyration with a `SIGMA` of (0.01,0.006) nm respectively. The simulations whose results are shown in Fig. S15(b,c), are again analogous to the OneOPES in the main text, with the difference that the `BARRIER` of the main OPES Explore is 100 kJ/mol and the `SIGMA` is determined adaptively.

The results in the main text are compared to the simulation from the seminal paper in Ref. [16] which is over 100 $\mu$s long.

For the thermodynamic properties evaluation, we took replica **6** of each OneOPES simulation and, after estimating $\Delta F$ between 340K and 360K through reweighting [19], we performed a linear fit of the Van 't Hoff equation $\Delta F = \Delta H - T\Delta S$ (see Fig. S19). In Table S4 we report the $R^2$ parameter of each fit and the slope and intercept of the linear fit that correspond to $-\Delta S$ and $\Delta U$, respectively. In the main text we report the average and the standard deviation of their values. By intersecting the linear fits with $\Delta F = 0$, it is trivial to estimate the protein melting temperature $T_{\mathrm{m}}$. Performing the same analysis on other replicas that include OPES MultiThermal gave analogous results.

#### Computational Performance

Running a OneOPES 8-replica simulation on a single modern workstation with an AMD 7950X CPU with 16 physical cores and an nVidia RTX 4090, we measured a performance of $\approx$ 10000 ns/day in Alanine Dipeptide, of $\approx$ 75 ns/day in Trypsin-Benzamidine and of $\approx$ 150 ns/day in Chignolin. Therefore, to reach the simulation length presented in the paper, one needs $\approx$ 7 minutes for Alanine Dipeptide, $\approx$ 3.3 days for Trypsin-Benzamidine and $\approx$ 2.7 days for Chignolin.

### Supporting Figures and Tables

#### Alanine Dipeptide

In the following sections, we report all the supplementary data we generated to support our investigation on the Alanine Dipeptide.

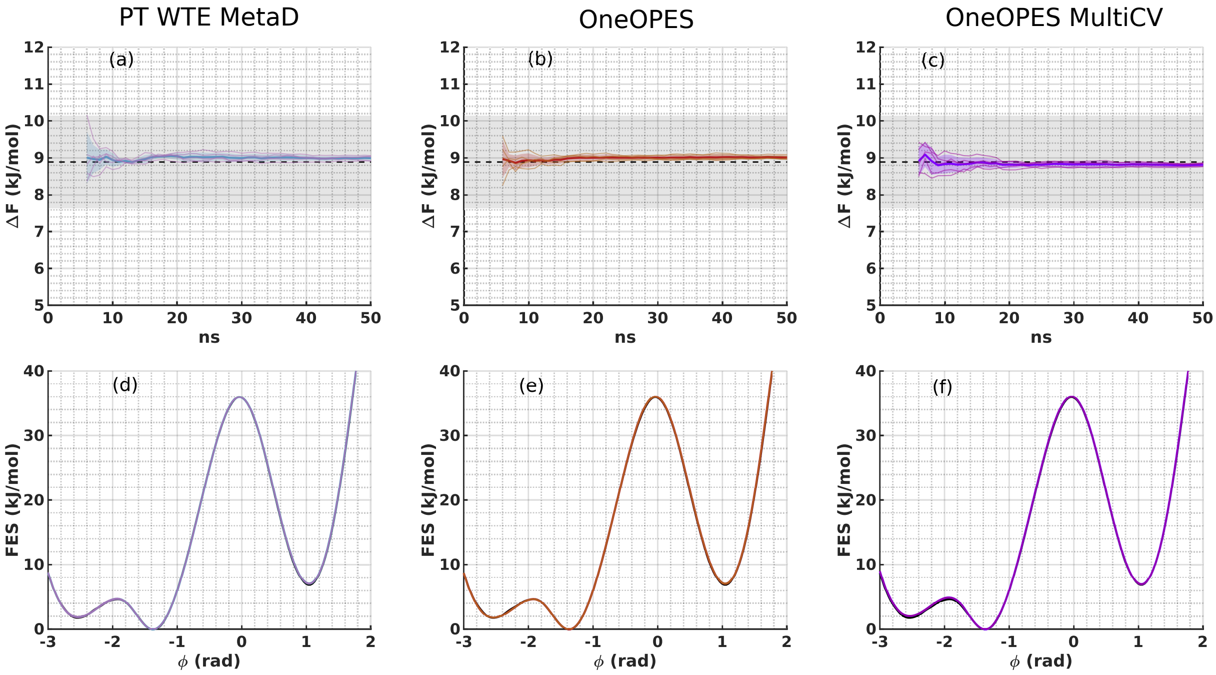


Figure S1: Set of 5 independent simulations on the Alanine Dipeptide system, where we bias the ideal CV $\phi$ with PT-WTE-MetaD (panels (a) and (c)), and OneOPES (panels (b) and (d)), and OneOPES MultiCV (panels (b) and (d)). In (a), (b), and (c) we show the average $\Delta F$ in time through dark blue, dark red, and dark purple solid lines, respectively. Similarly, their standard deviation is displayed through the semitransparent regions and coloured in light blue, light red, and light purple. $\Delta F$ values corresponding to individual OneOPES simulations are shown in solid purple, orange, and magenta lines, respectively. The expected $\Delta F$ is indicated by a dashed black line with an error of 0.5 $k_B T$ in shaded grey. In (d), (e), and (f), we show the one-dimensional FES reweighted over $\phi$ after 40 ns for PT-WTE-MetaD, OneOPES, and OneOPES MultiCV simulations, respectively. The same colour scheme applies as in panels (a), (b), and (c).

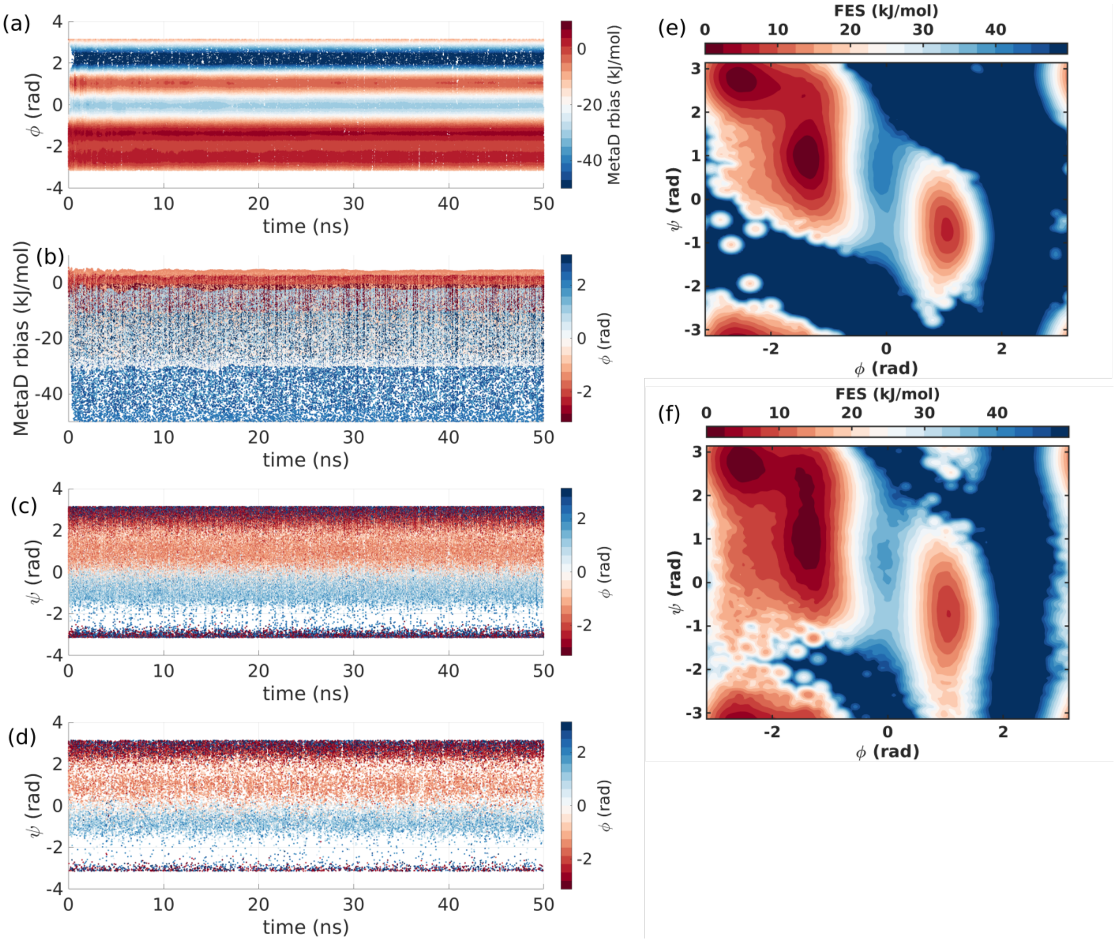


Figure S2: A representative PT-WTE-MetaD simulation of Alanine Dipeptide in which we bias the optimal CV $\phi$. In (a), we display the dynamics of the CV $\phi$, coloured based upon the deposited MetaD *rbias*. In (b), we show the dynamics of the MetaD *rbias* in replica **0**, coloured according to the values assumed by the CV $\phi$. In (c), we show the dynamics of the CV $\psi$ in replica **0**, coloured according to the values assumed by the CV $\phi$. In (d), we show the dynamics of the CV $\psi$ in the *demuxed* replica **0**, coloured according to the values assumed by the CV $\phi$. In (e) and (f), we show the 2D FES along the dihedral angles $\phi$ and $\psi$, obtained through reweighting at the end of replica **0** and **7**, respectively.

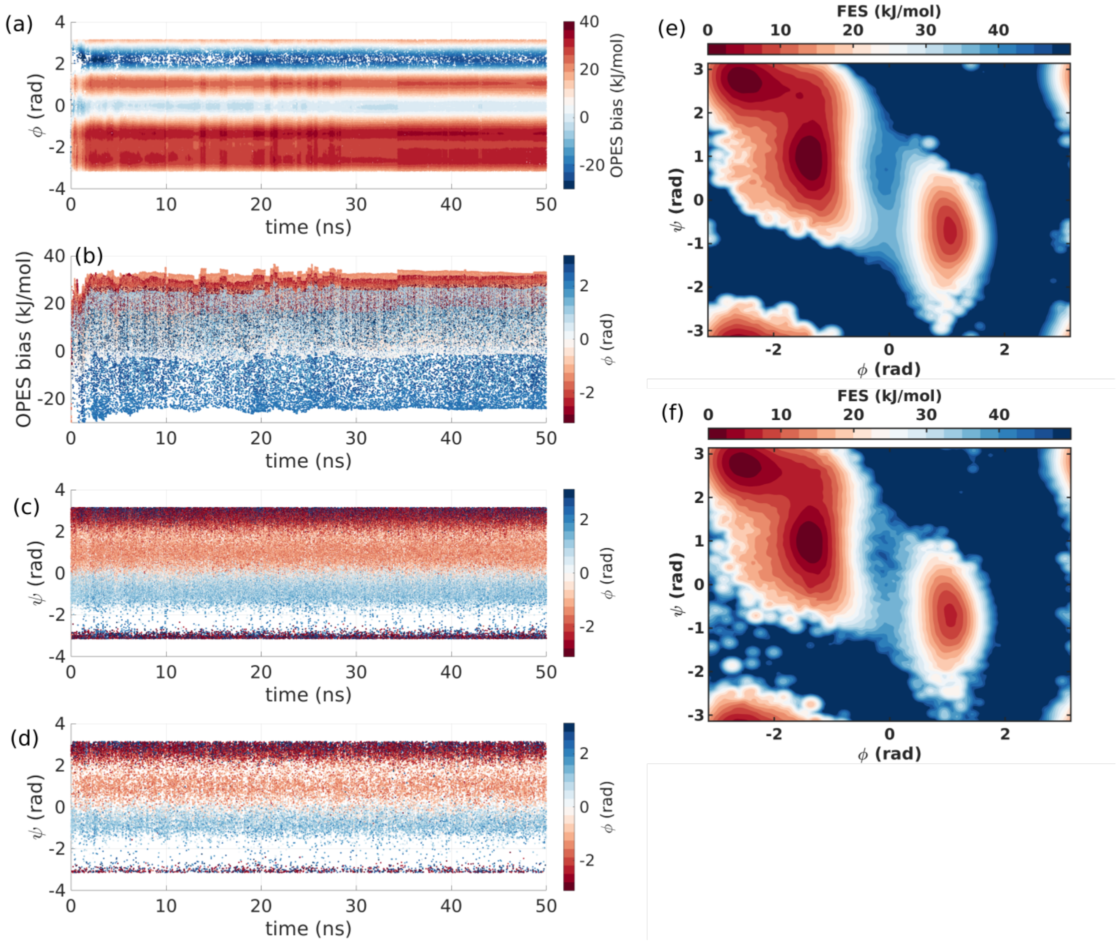


Figure S3: A representative OneOPES simulation of Alanine Dipeptide in which we bias the optimal CV $\phi$. In (a), we display the dynamics of the CV $\phi$, coloured based on the OPES bias. In (b), we show the dynamics of OPES bias in replica **0**, coloured according to the values assumed by the CV $\phi$. In (c), we show the dynamics of the CV $\psi$ in replica **0**, coloured according to the values assumed by the CV $\phi$. In (d), we show the dynamics of the CV $\psi$ in the *demuxed* replica **0**, coloured according to the values assumed by the CV $\phi$. In (e) and (f), we show the 2D FES along the dihedral angles $\phi$ and $\psi$, obtained through reweighting at the end of replica **0** and **7**, respectively.

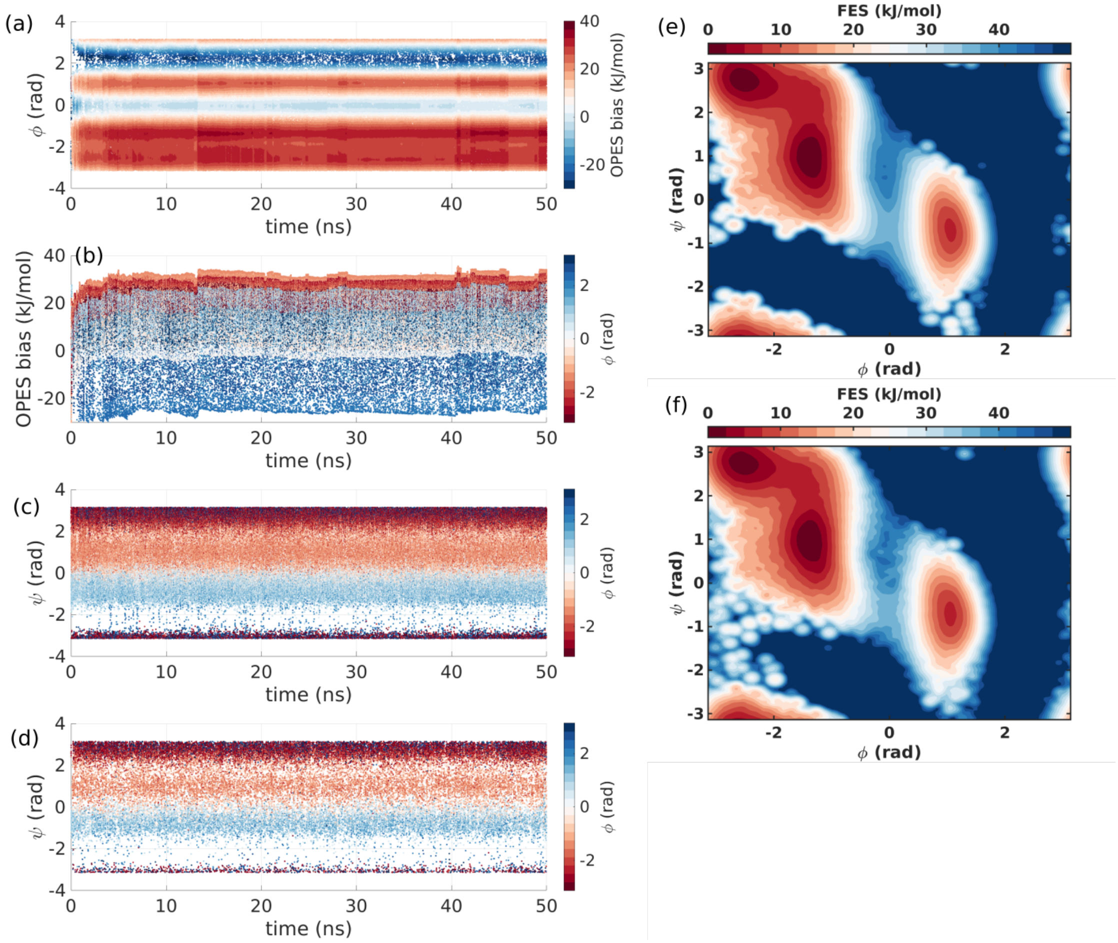


Figure S4: A representative OneOPES MultiCV simulation of Alanine Dipeptide in which we bias the optimal CV $\phi$. In (a), we display the dynamics of the CV $\phi$, coloured based on the OPES bias. In (b), we show the dynamics of the OPES bias in replica **0**, coloured according to the values assumed by the CV $\phi$. In (c), we show the dynamics of the CV $\psi$ in replica **0**, coloured according to the values assumed by the CV $\phi$. In (d), we show the dynamics of the CV $\psi$ in the *demuxed* replica **0**, coloured according to the values assumed by the CV $\phi$. In (e) and (f), we show the 2D FES along the dihedral angles $\phi$ and $\psi$, obtained through reweighting at the end of replica **0** and **7**, respectively.

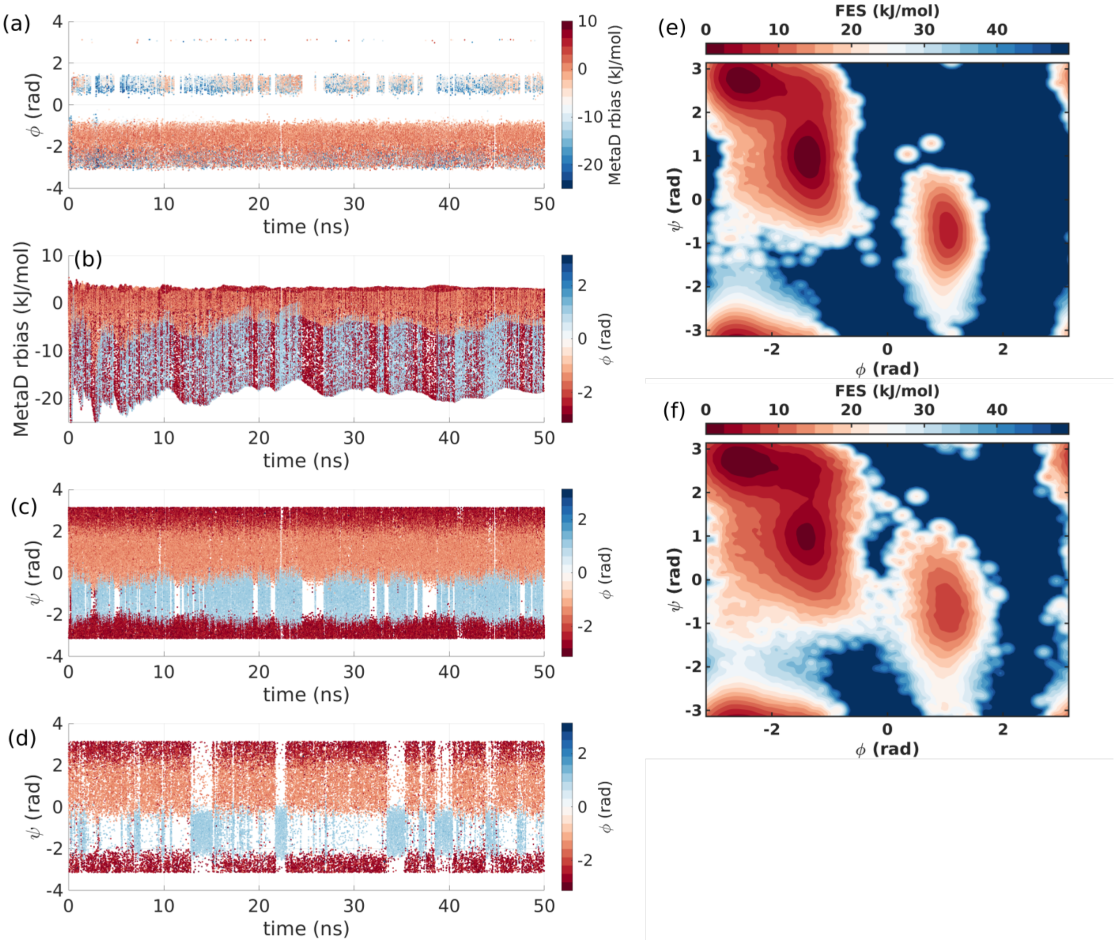


Figure S5: A representative PT-WTE-MetaD simulation of Alanine Dipeptide in which we bias the sub-optimal CV $\psi$. In (a), we display the dynamics of the CV $\phi$, coloured based upon the deposited MetaD *rbias*. In (b), we show the dynamics of the MetaD *rbias* in replica **0**, coloured according to the values assumed by the CV $\phi$. In (c), we show the dynamics of the CV $\psi$ in replica **0**, coloured according to the values assumed by the CV $\phi$. In (d), we show the dynamics of the CV $\psi$ in the *demuxed* replica **0**, coloured according to the values assumed by the CV $\phi$. In (e) and (f), we show the 2D FES along the dihedral angles $\phi$ and $\psi$, obtained through reweighting at the end of replica **0** and **7**, respectively.

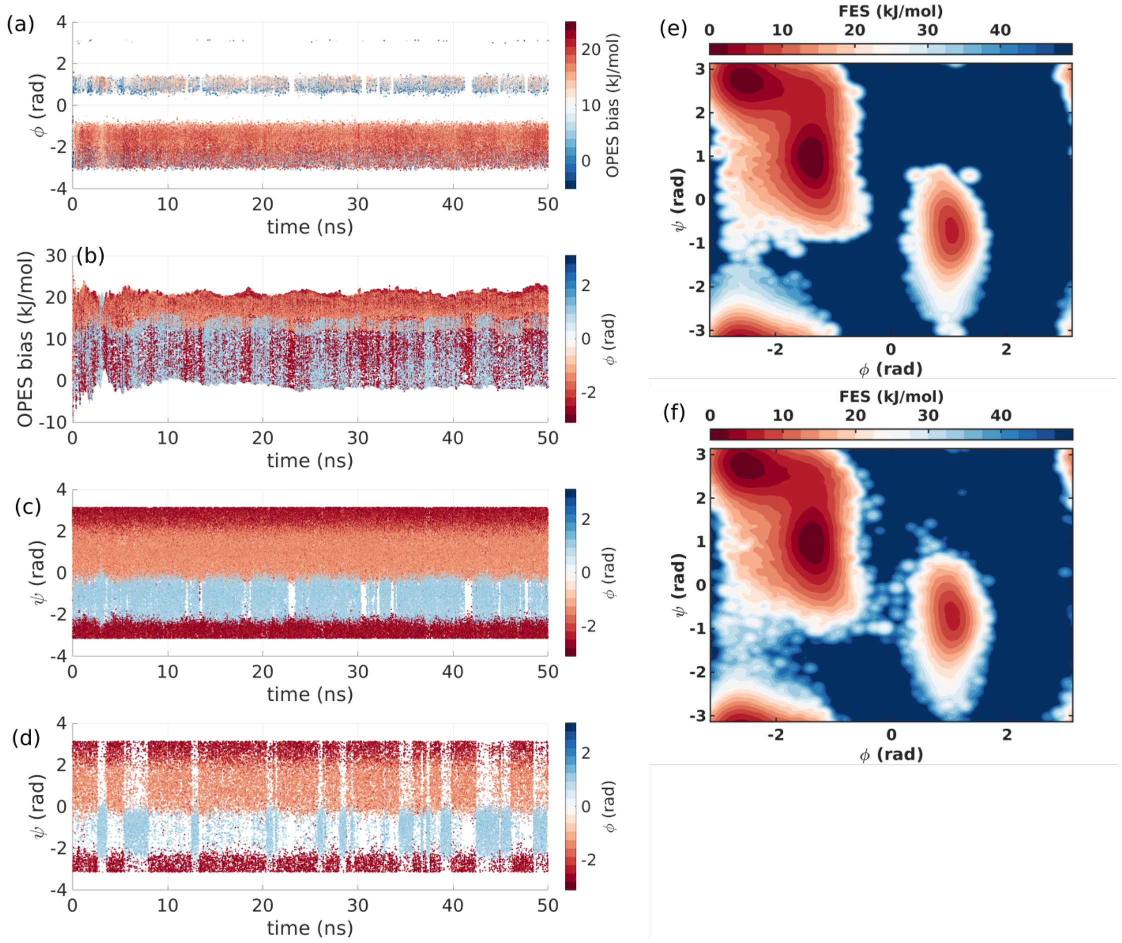


Figure S6: A representative OneOPES simulation of Alanine Dipeptide in which we bias the sub-optimal CV $\psi$. In (a), we display the dynamics of the CV $\phi$, coloured based on the OPES bias. In (b), we show the dynamics of the OPES bias in replica **0**, coloured according to the values assumed by the CV $\phi$. In (c), we show the dynamics of the CV $\psi$ in replica **0**, coloured according to the values assumed by the CV $\phi$. In (d), we show the dynamics of the CV $\psi$ in the *demuxed* replica **0**, coloured according to the values assumed by the CV $\phi$. In (e) and (f), we show the 2D FES along the dihedral angles $\phi$ and $\psi$, obtained through reweighting at the end of replica **0** and **7**, respectively.

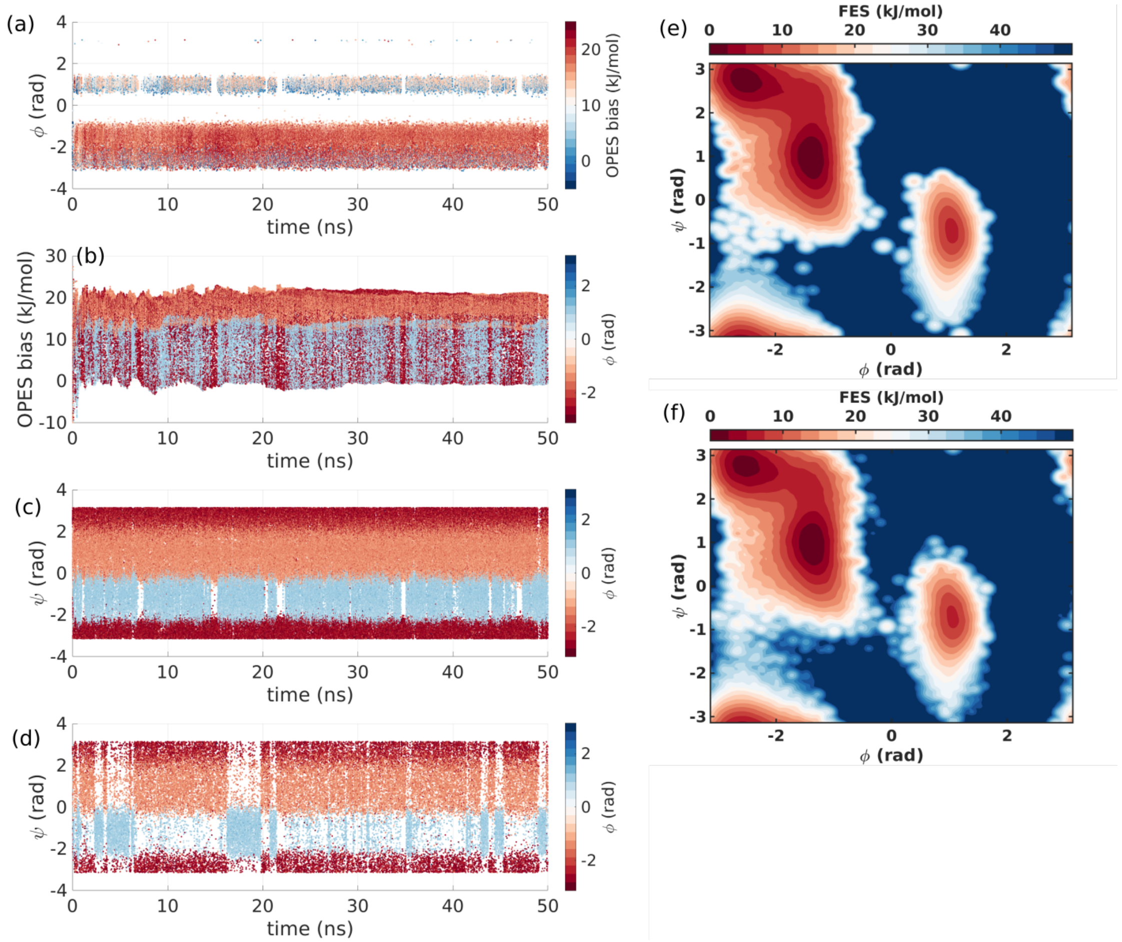


Figure S7: A representative OneOPES MultiCV simulation of Alanine Dipeptide in which we bias the sub-optimal CV $\psi$. In (a), we display the dynamics of the CV $\phi$, coloured based on the OPES bias. In (b), we show the dynamics of the OPES bias in replica **0**, coloured according to the values assumed by the CV $\phi$. In (c), we show the dynamics of the CV $\psi$ in replica **0**, coloured according to the values assumed by the CV $\phi$. In (d), we show the dynamics of the CV $\psi$ in the *demuxed* replica **0**, coloured according to the values assumed by the CV $\phi$. In (e) and (f), we show the 2D FES along the dihedral angles $\phi$ and $\psi$, obtained through reweighting at the end of replica **0** and **7**, respectively.

Table S1: Average replica exchange probabilities collected on the Alanine Dipeptide PT-WTE-MetaD, OneOPES, and OneOPES MultiCV simulations.

| Method | CV | R0-R1 | R1-R2 | R2-R3 | R3-R4 | R4-R5 | R5-R6 | R6-R7 |
|---|---|---|---|---|---|---|---|---|
| PT-WTE-MetaD | $\phi$ | 0.92 | 0.92 | 0.92 | 0.16 | 0.66 | 0.62 | 0.67 |
| OneOPES | $\phi$ | 0.82 | 0.82 | 0.82 | 0.34 | 0.76 | 0.79 | 0.82 |
| OneOPES MultiCV | $\phi$ | 0.82 | 0.80 | 0.81 | 0.34 | 0.81 | 0.80 | 0.76 |
| PT-WTE-MetaD | $\psi$ | 0.93 | 0.93 | 0.93 | 0.23 | 0.74 | 0.70 | 0.72 |
| OneOPES | $\psi$ | 0.80 | 0.79 | 0.80 | 0.37 | 0.80 | 0.79 | 0.79 |
| OneOPES MultiCV | $\psi$ | 0.80 | 0.78 | 0.78 | 0.38 | 0.76 | 0.78 | 0.79 |

#### Trypsin-Benzamidine

In the following sections, we report all the supplementary data we generated to support our investigation on the Trypsine-Benzamidine binding process.

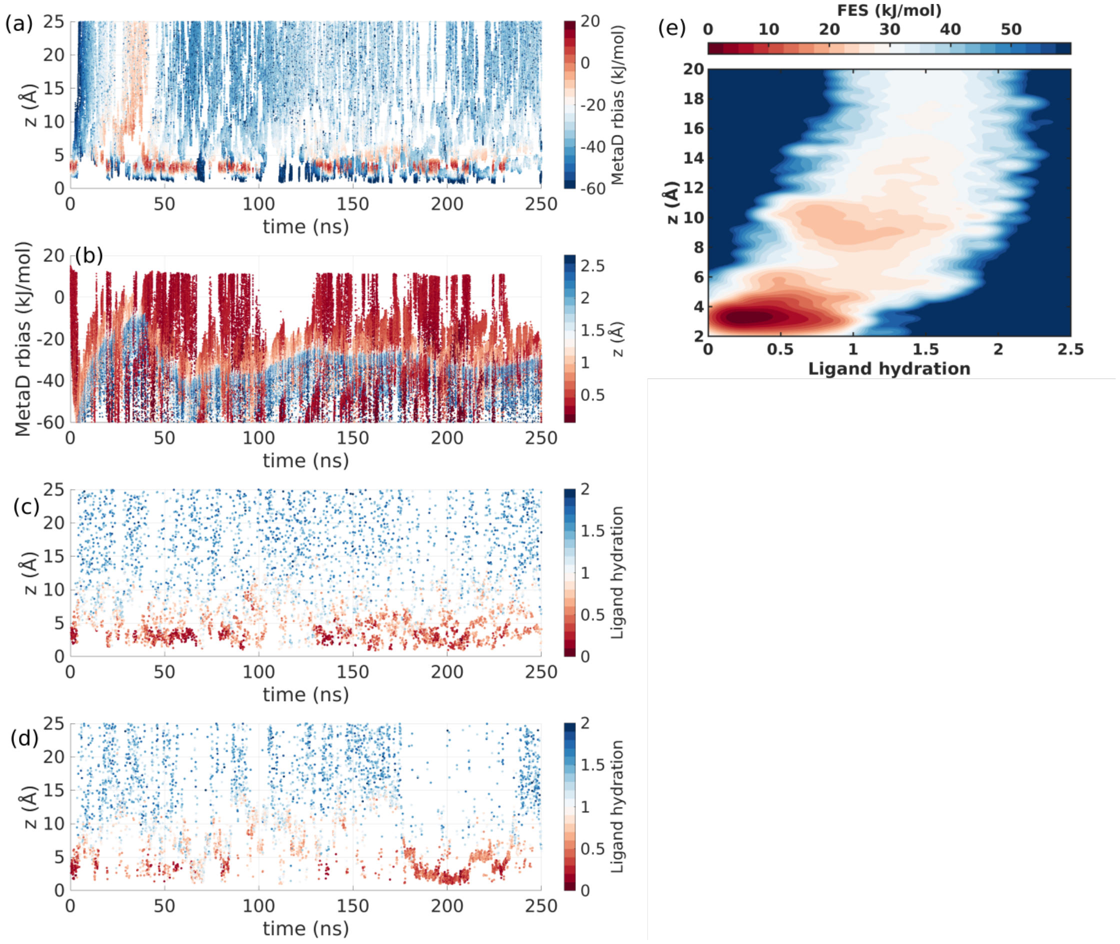


Figure S8: A representative PT-WTE-MetaD simulation of the Trypsin-Benzamidine system, in which we bias the Funnel's CVs $z$ and $r$. In (a), we display the dynamics of the CV $z$, coloured based upon the deposited MetaD *rbias*. In (b), we show the dynamics of the MetaD *rbias* in replica **0**, coloured according to the values assumed by the CV $z$. In (c), we show the dynamics of the CV $z$ in replica **0**, coloured according to the values assumed by the CV **G** (named "*Ligand hydration*" hereafter). In (d), we show the dynamics of the CV $z$ in the *demuxed* replica **0**, coloured according to the values assumed by the CV *Ligand hydration*. In (e), we show the 2D FES along the CVs $z$ and *Ligand hydration*, obtained through reweighting on replica **0** at the end of the PT-WTE-MetaD simulation.

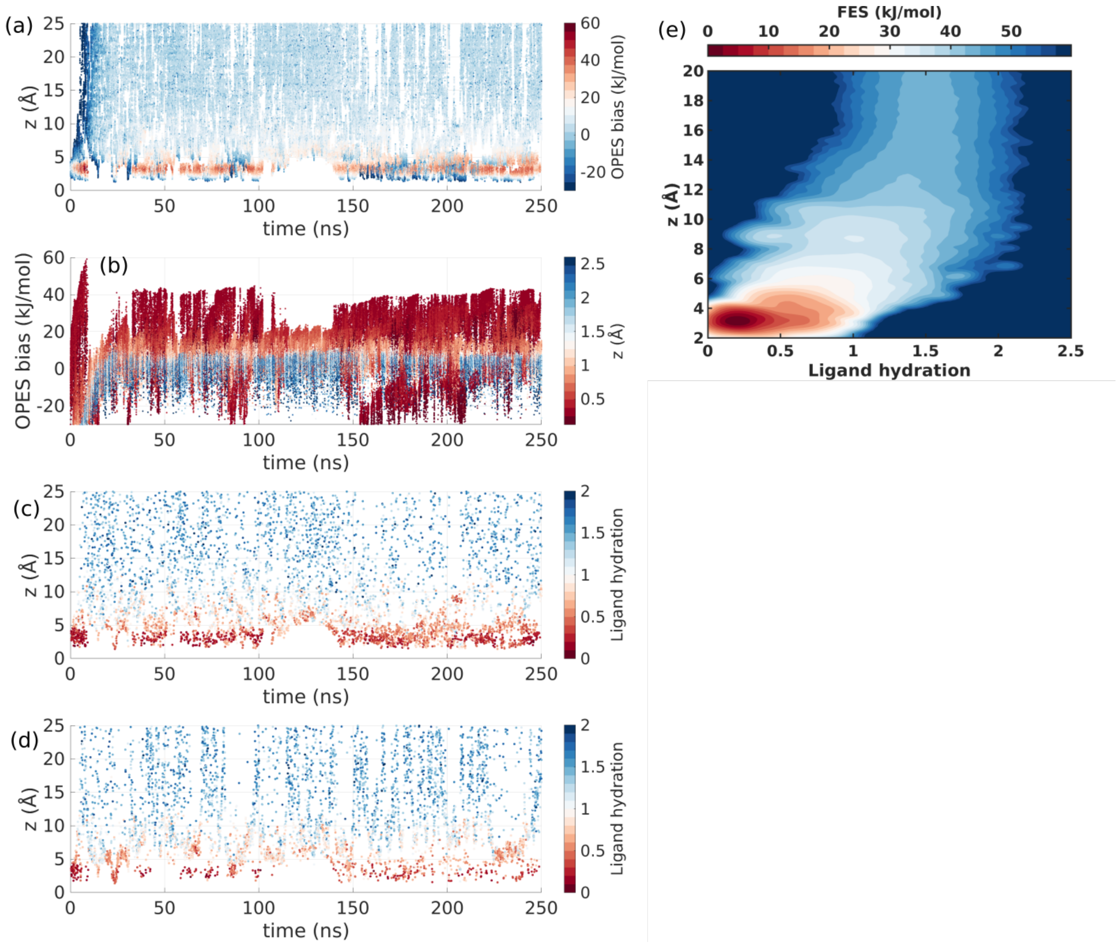


Figure S9: A representative OneOPES simulation of the Trypsin-Benzamidine system, in which we bias the Funnel's CVs $z$ and $r$. In (a), we display the dynamics of the CV $z$, coloured based on the OPES bias. In (b), we show the dynamics of the OPES bias in replica **0**, coloured according to the values assumed by the CV $z$. In (c), we show the dynamics of the CV $z$ in replica **0**, coloured according to the values assumed by the CV **G** (named "*Ligand hydration*" hereafter). In (d), we show the dynamics of the CV $z$ in the *demuxed* replica **0**, coloured according to the values assumed by the CV *Ligand hydration*. In (e), we show the 2D FES along the CVs $z$ and *Ligand hydration*, obtained through reweighting on replica **0** at the end of the OneOPES simulation.

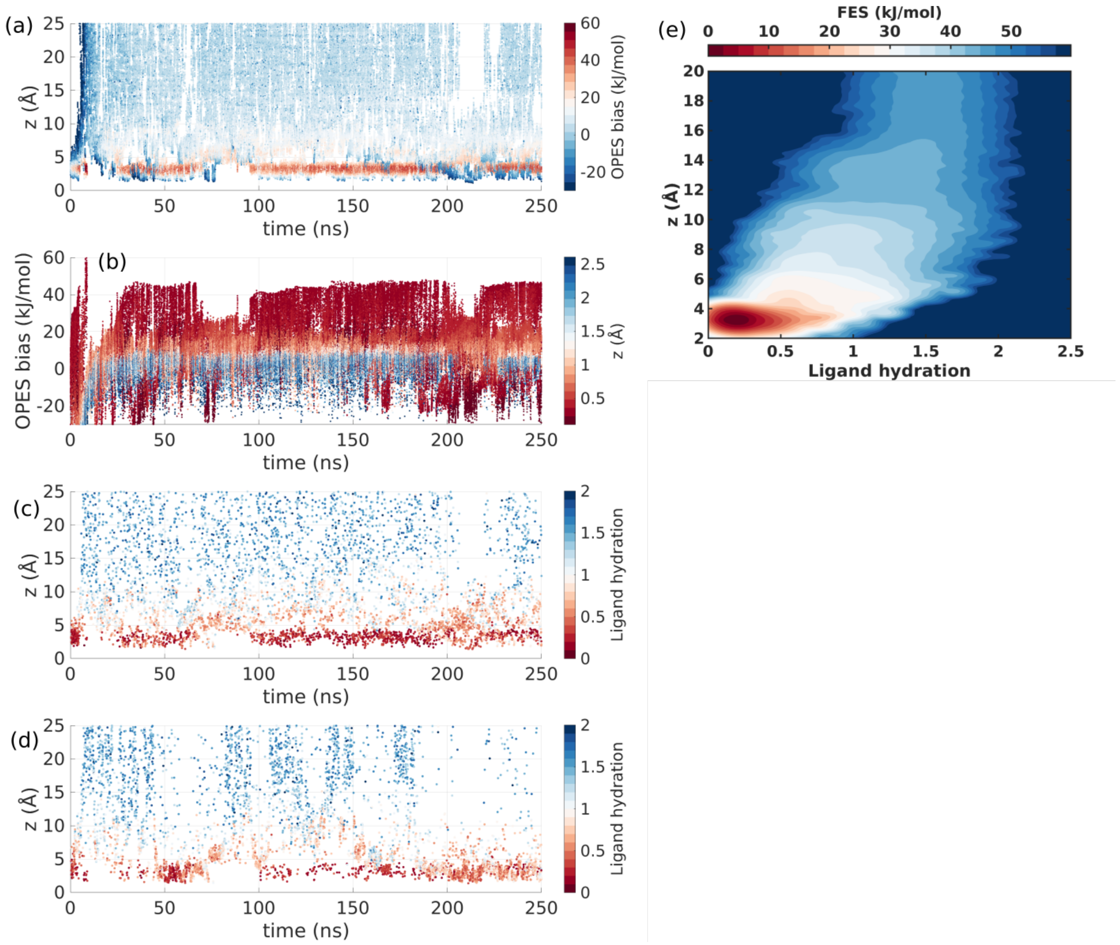


Figure S10: A representative OneOPES MultiCV simulation of the Trypsin-Benzamidine system, in which we bias the Funnel's CVs $z$ and $r$. In (a), we display the dynamics of the CV $z$, coloured based on the OPES bias. In (b), we show the dynamics of the OPES bias in replica **0**, coloured according to the values assumed by the CV $z$. In (c), we show the dynamics of the CV $z$ in replica **0**, coloured according to the values assumed by the CV **G** (named "*Ligand hydration*" hereafter). In (d), we show the dynamics of the CV $z$ in the *demuxed* replica **0**, coloured according to the values assumed by the CV *Ligand hydration*. In (e), we show the 2D FES along the CVs $z$ and *Ligand hydration*, obtained through reweighting on replica **0** at the end of the OneOPES MultiCV simulation.

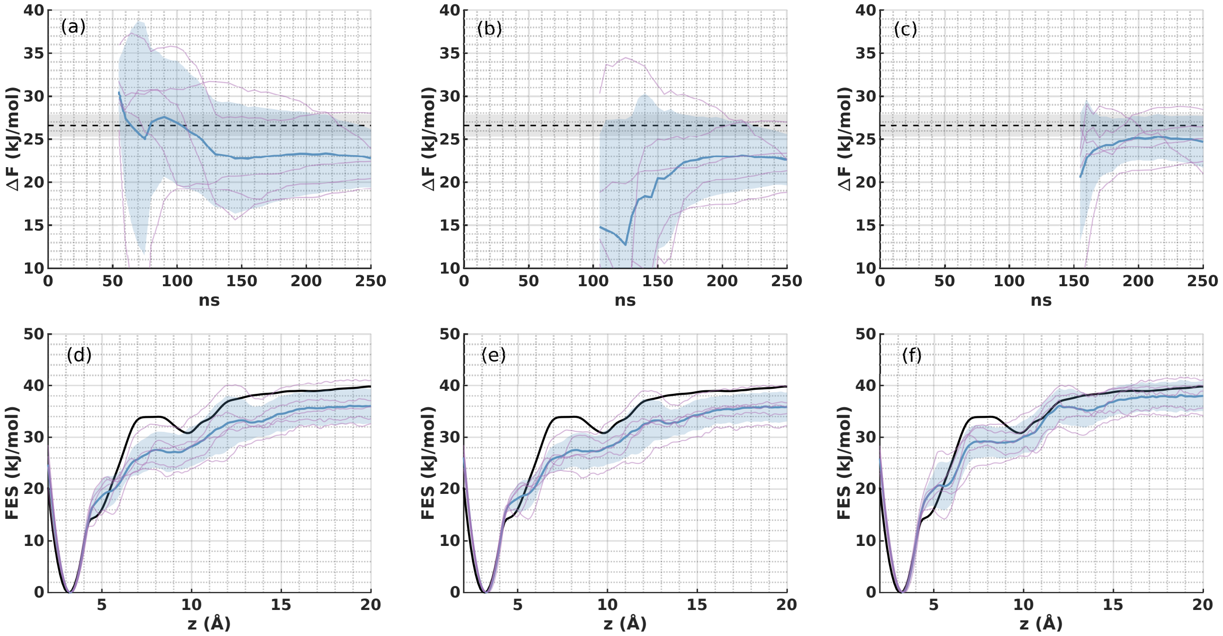


Figure S11: Set of 5 independent PT-WTE-MetaD simulations on the Trypsin-Benzamidine system, where we bias the Funnel's CVs $z$ and $r$. Herein, we report the effect of discarding more of the initial portion of the data on the final estimate of the binding free energy. In panels (a), (b), and (c), we show the average $\Delta F$ in time obtained by repeating the reweighting procedure and discarding respectively 20%, 40%, and 60% of the replica **0** trajectories. By following the same protocol, we reweight the one-dimensional FES on the CV $z$, displayed in panels (d), (e), and (f), respectively. We show the average $\Delta F$ in time through blue solid lines. The standard deviation is displayed through semitransparent regions in light blue. $\Delta F$ values corresponding to individual OneOPES simulations are shown in solid magenta lines. The expected $\Delta F$ is indicated by a dashed black line with an error of 0.5 $k_{\mathrm{B}}T$ in shaded grey.

Table S2: Average replica exchange probabilities collected on the Trypsin-Benzamidine PT-WTE-MetaD, OneOPES, and OneOPES MultiCV simulations.

| Method | R0-R1 | R1-R2 | R2-R3 | R3-R4 | R4-R5 | R5-R6 | R6-R7 |
|---|---|---|---|---|---|---|---|
| PT-WTE-MetaD | 0.83 | 0.86 | 0.77 | 0.10 | 0.21 | 0.14 | 0.10 |
| OneOPES | 0.57 | 0.56 | 0.57 | 0.15 | 0.27 | 0.39 | 0.37 |
| OneOPES MultiCV | 0.56 | 0.55 | 0.57 | 0.15 | 0.31 | 0.33 | 0.36 |

#### Chignolin

In the following sections, we report all the supplementary data we generated to support our investigation on the folding of Chignolin.

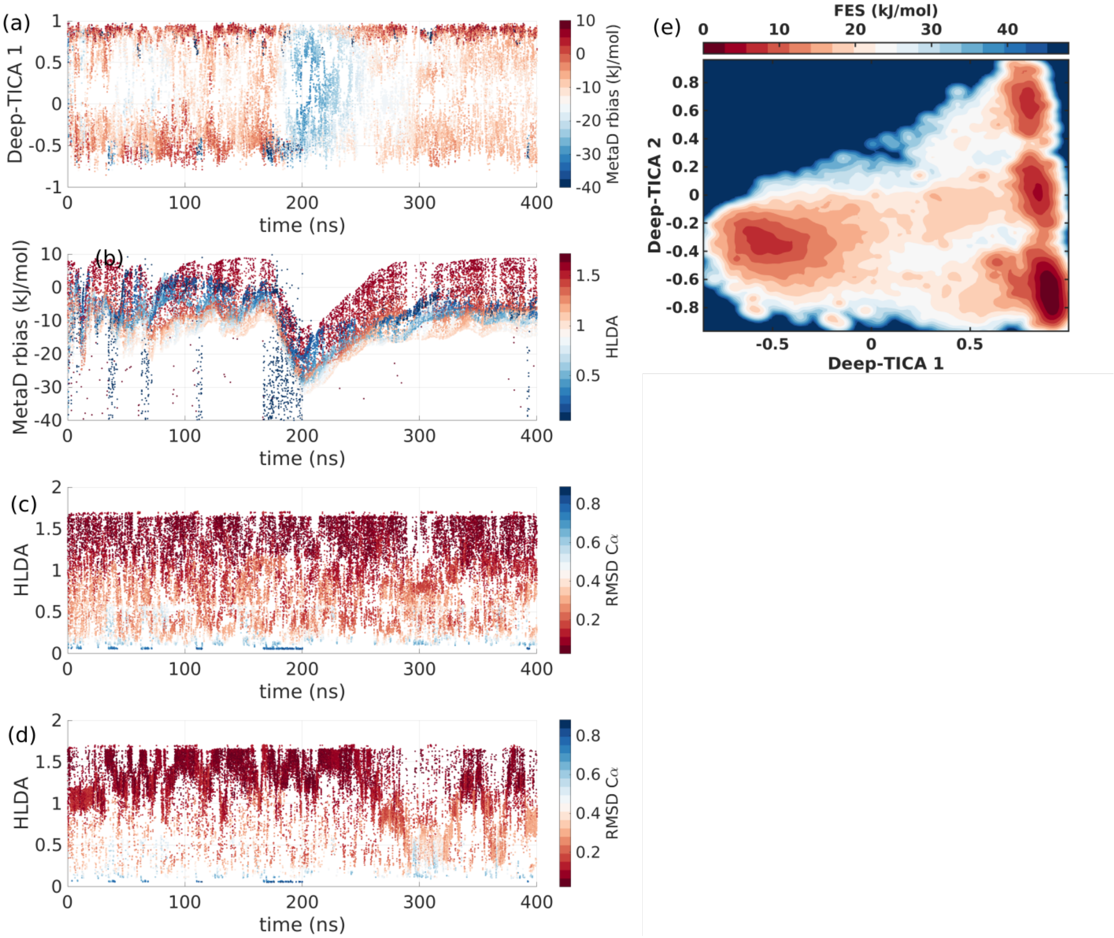


Figure S12: A representative PT-WTE-MetaD simulation of the Chignolin system, in which we bias the CVs *HLDA*. In (a), we display the dynamics of the ideal *Deep-TICA1* CV from Ref. [20], coloured based upon the deposited MetaD *rbias*. In (b), we show the dynamics of the MetaD *rbias* in replica **0**, coloured according to the values assumed by the CV *HLDA*. In (c), we show the dynamics of the CV *HLDA* in replica **0**, coloured according to the values assumed by the CV *RMSD* (see "Supplementary Computational Details"). In (d), we show the dynamics of the CV *HLDA* in the *demuxed* replica **0**, coloured according to the values assumed by the CV *RMSD*. In (e), we show the 2D FES along the ideal CVs *Deep-TICA1* and *Deep-TICA2* from Ref. [20], obtained through reweighting on replica **0** at the end of the PT-WTE-MetaD simulation.

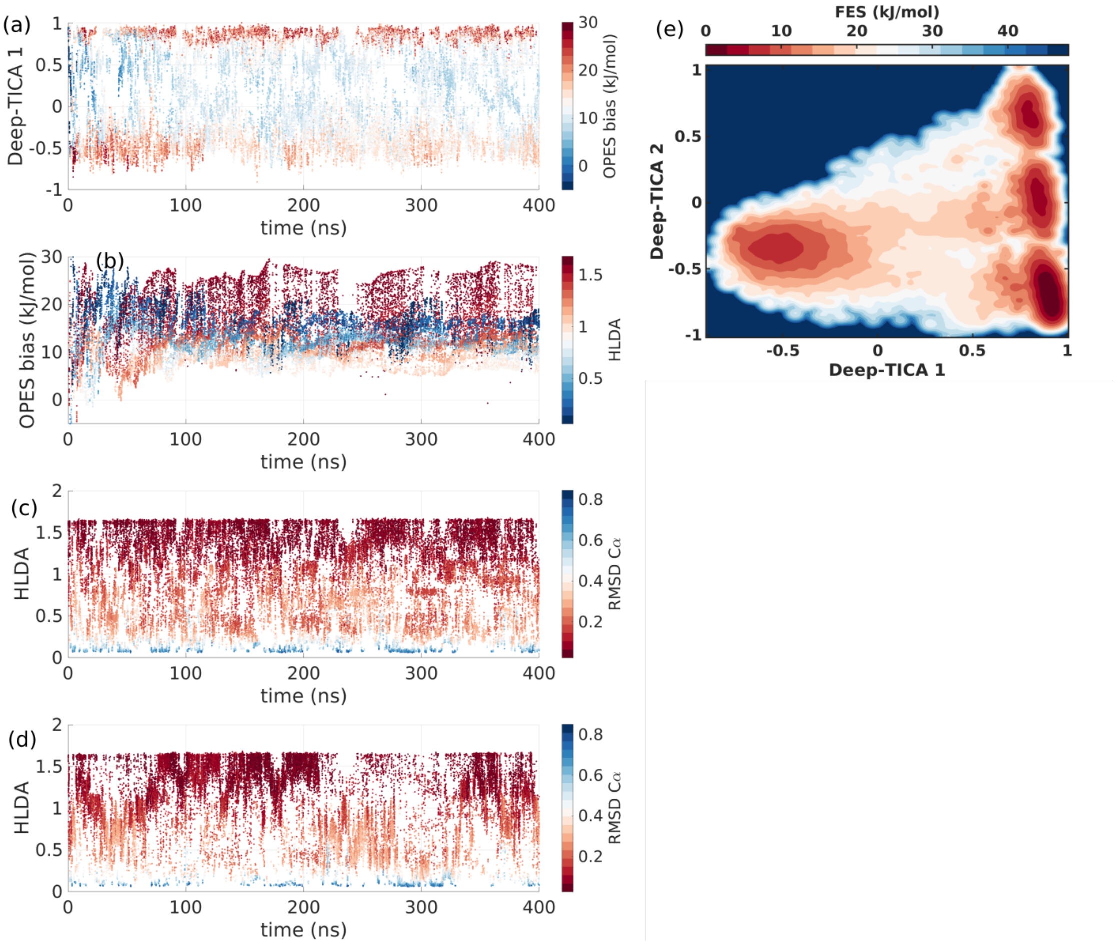


Figure S13: A representative OneOPES simulation of the Chignolin system, in which we bias the CVs *HLDA*. In (a), we display the dynamics of the ideal *Deep-TICA1* CV from Ref. [20], coloured based on the OPES bias. In (b), we show the dynamics of the OPES bias in replica **0**, coloured according to the values assumed by the CV *HLDA*. In (c), we show the dynamics of the CV *HLDA* in replica **0**, coloured according to the values assumed by the CV *RMSD* (see "Supplementary Computational Details"). In (d), we show the dynamics of the CV *HLDA* in the *demuxed* replica **0**, coloured according to the values assumed by the CV *RMSD*. In (e), we show the 2D FES along the ideal CVs *Deep-TICA1* and *Deep-TICA2* from Ref. [20], obtained through reweighting on replica **0** at the end of the OneOPES simulation.

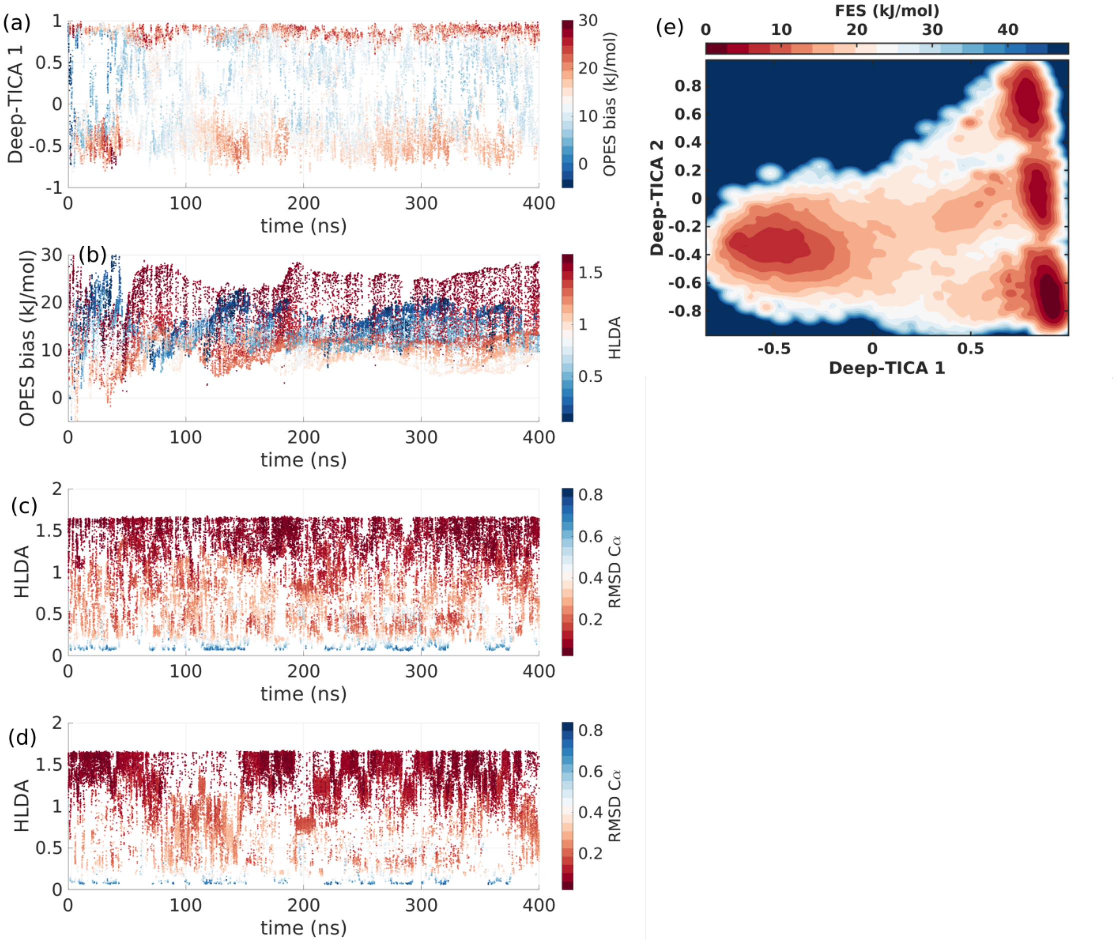


Figure S14: A representative OneOPES MultiCV simulation of the Chignolin system, in which we bias the CVs *HLDA*. In (a), we display the dynamics of the ideal *Deep-TICA1* CV from Ref. [20], coloured based on the OPES bias. In (b), we show the dynamics of the OPES bias in replica **0**, coloured according to the values assumed by the CV *HLDA*. In (c), we show the dynamics of the CV *HLDA* in replica **0**, coloured according to the values assumed by the CV *RMSD* (see "Supplementary Computational Details"). In (d), we show the dynamics of the CV *HLDA* in the *demuxed* replica **0**, coloured according to the values assumed by the CV *RMSD*. In (e), we show the 2D FES along the ideal CVs *Deep-TICA1* and *Deep-TICA2* from Ref. [20], obtained through reweighting on replica **0** at the end of the OneOPES MultiCV simulation.

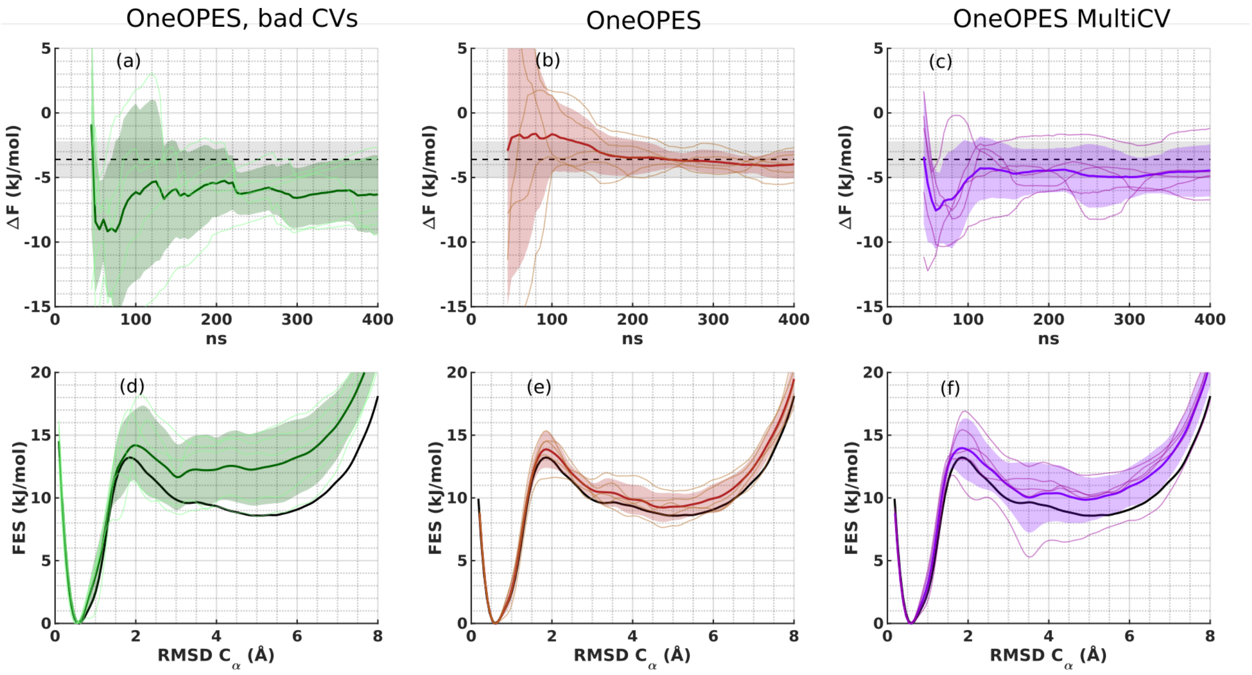


Figure S15: Set of 5 independent simulations on the Chignolin system, where we carried out OneOPES and OneOPES MultiCV simulations employing different sampling parameters. Panels (a) and (d) show the outcomes of running OneOPES on the "bad" CVs *RMSD* and *rg*, i.e. the average $\Delta F$ in time and the one-dimensional FES reweighted over the *RMSD* CV, respectively. The average $\Delta F$ is coloured in dark green, while its standard deviation is displayed through a semitransparent region and coloured in light green. $\Delta F$ values corresponding to individual OneOPES simulations are shown in solid green. Panels (b) and (e) show the outcomes of running OneOPES with a $\Delta E$=100 kJ/mol on the CV *HLDA*. The average $\Delta F$ is coloured in dark red, while its standard deviation is displayed through a semitransparent light red region. $\Delta F$ values corresponding to individual OneOPES simulations are shown in solid red. Panels (c) and (f) show the outcomes of running OneOPES MultiCV with a $\Delta E$=100 kJ/mol on the CV *HLDA* (and the auxiliary variables described in "Supplementary Computational Details"). The average $\Delta F$ is coloured in dark purple, while its standard deviation is displayed through a semitransparent light purple region. $\Delta F$ values corresponding to individual OneOPES simulations are shown in solid magenta. In all panels, the expected $\Delta F$ is indicated by a dashed black line with an error of 0.5 $k_\mathrm{B}T$ in shaded grey.

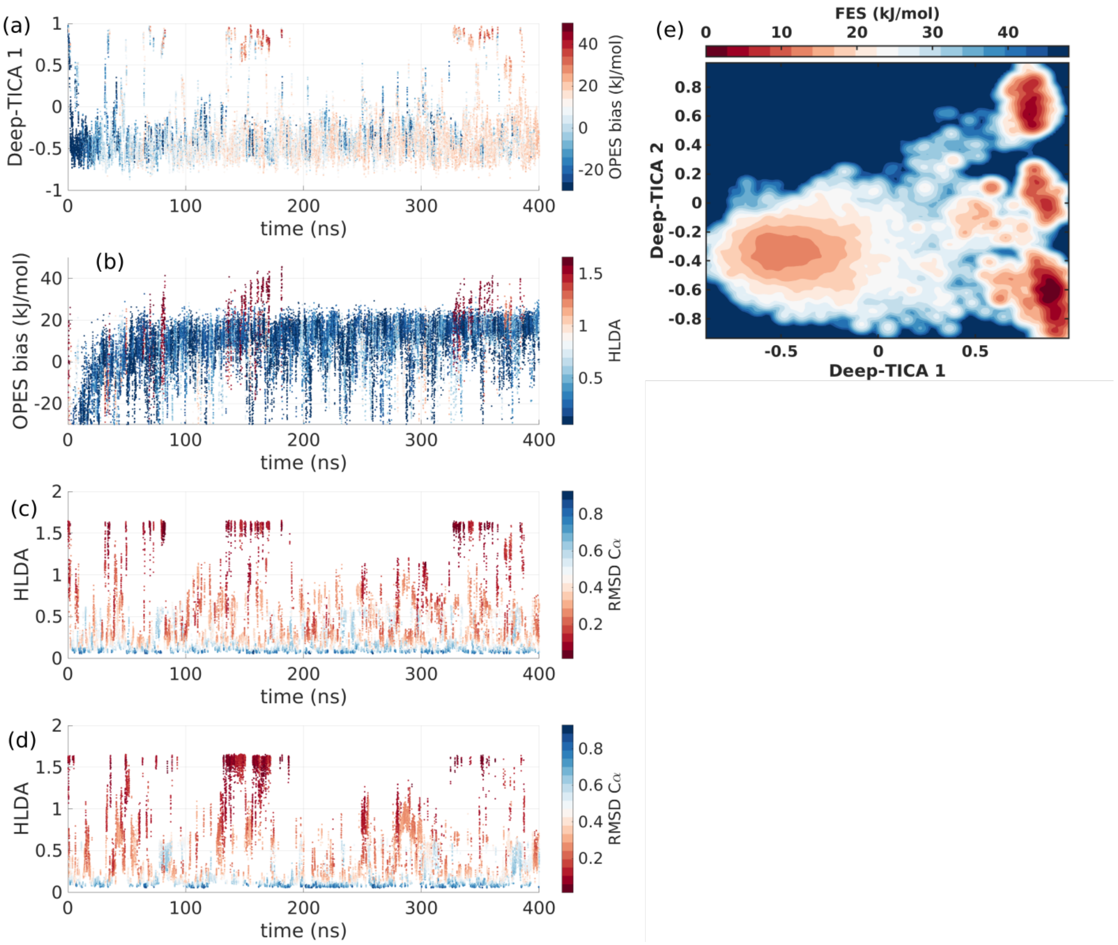


Figure S16: A representative OneOPES simulation of the Chignolin system, in which we bias the "bad" CVs *RMSD* and *rg*. In (a), we display the dynamics of the ideal *Deep-TICA1* CV from Ref. [20], coloured based on the OPES bias. In (b), we show the dynamics of the OPES bias in replica **0**, coloured according to the values assumed by the CV *HLDA*. In (c), we show the dynamics of the CV *HLDA* in replica **0**, coloured according to the values assumed by the CV *RMSD*. In (d), we show the dynamics of the CV *HLDA* in the *demuxed* replica **0**, coloured according to the values assumed by the CV *RMSD*. In (e), we show the 2D FES along the ideal CVs *Deep-TICA1* and *Deep-TICA2* from Ref. [20], obtained through reweighting on replica **0** at the end of the OneOPES MultiCV simulation.

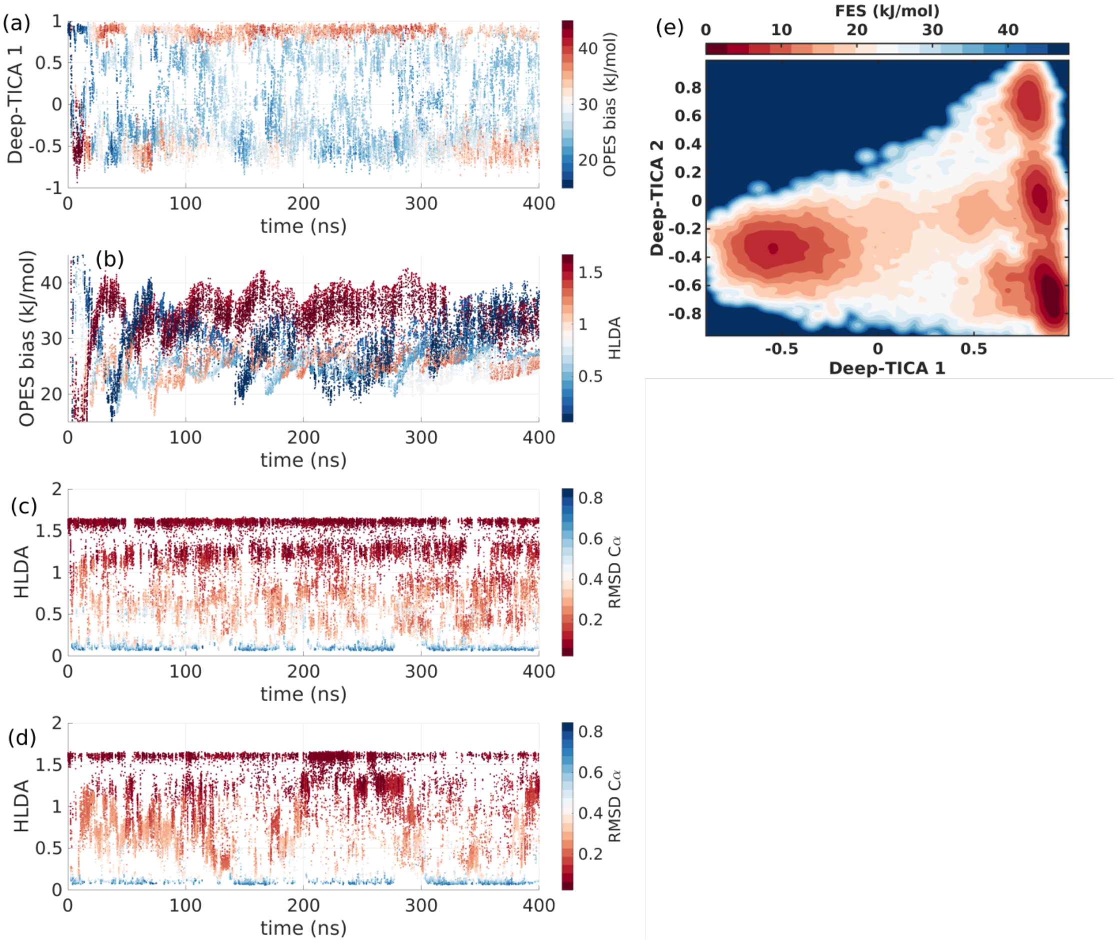


Figure S17: A representative OneOPES simulation of the Chignolin system, in which we bias the CV *HLDA* employing a $\Delta E$ of 100 kJ/mol. In (a), we display the dynamics of the ideal *Deep-TICA1* CV from Ref. [20], coloured based on the OPES bias. In (b), we show the dynamics of the OPES bias in replica **0**, coloured according to the values assumed by the CV *HLDA*. In (c), we show the dynamics of the CV *HLDA* in replica **0**, coloured according to the values assumed by the CV *RMSD* (see "Supplementary Computational Details"). In (d), we show the dynamics of the CV *HLDA* in the *demuxed* replica **0**, coloured according to the values assumed by the CV *RMSD*. In (e), we show the 2D FES along the ideal CVs *Deep-TICA1* and *Deep-TICA2* from Ref. [20], obtained through reweighting on replica **0** at the end of the OneOPES simulation.

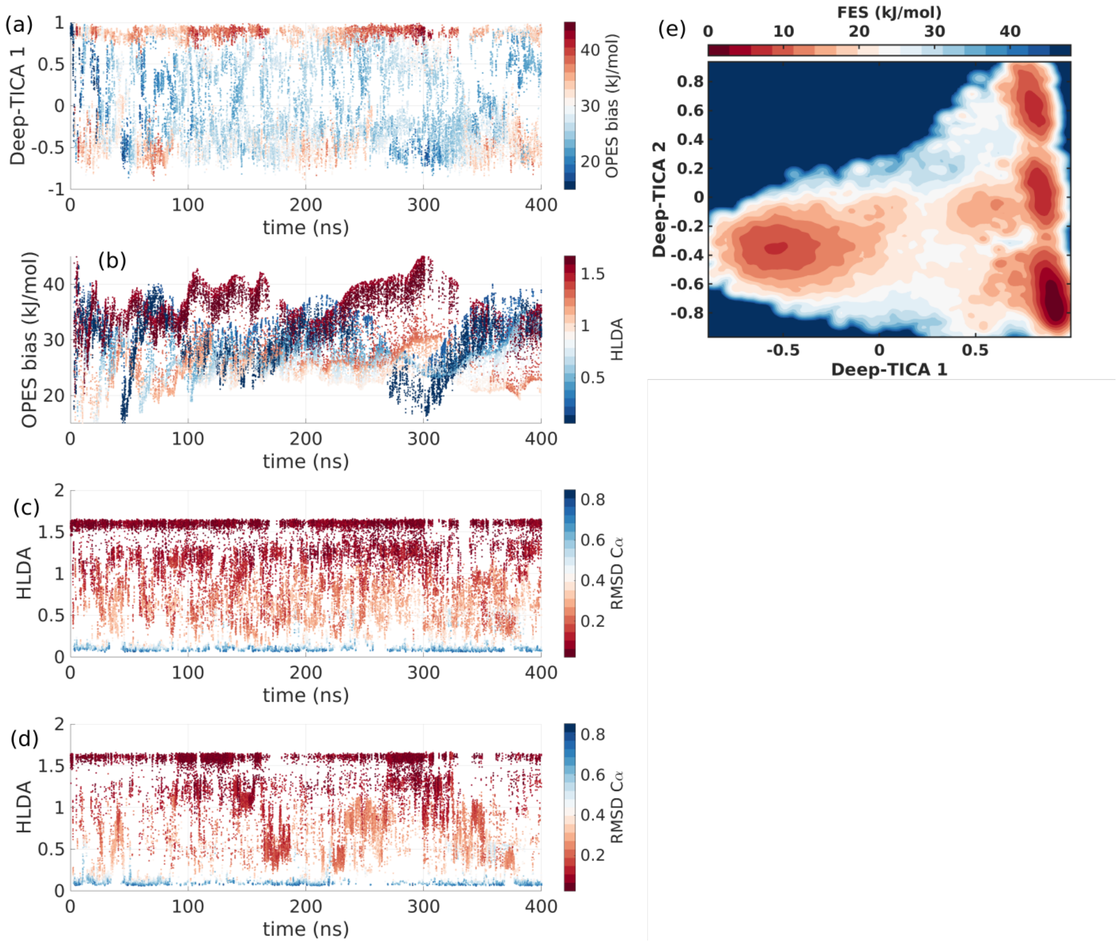


Figure S18: A representative OneOPES MultiCV simulation of the Chignolin system, in which we bias the CV *HLDA* employing a $\Delta E$ of 100 kJ/mol. In (a), we display the dynamics of the ideal *Deep-TICA1* CV from Ref. [20], coloured based on the OPES bias. In (b), we show the dynamics of the OPES bias in replica **0**, coloured according to the values assumed by the CV *HLDA*. In (c), we show the dynamics of the CV *HLDA* in replica **0**, coloured according to the values assumed by the CV *RMSD* (see "Supplementary Computational Details"). In (d), we show the dynamics of the CV *HLDA* in the *demuxed* replica **0**, coloured according to the values assumed by the CV *RMSD*. In (e), we show the 2D FES along the ideal CVs *Deep-TICA1* and *Deep-TICA2* from Ref. [20], obtained through reweighting on replica **0** at the end of the OneOPES MultiCV simulation.

Table S3: Average replica exchange probabilities collected on the Chignolin PT-WTE-MetaD, OneOPES, and OneOPES MultiCV simulations.

| Method | R0-R1 | R1-R2 | R2-R3 | R3-R4 | R4-R5 | R5-R6 | R6-R7 |
|---|---|---|---|---|---|---|---|
| PT-WTE-MetaD | 0.97 | 0.97 | 0.97 | 0.28 | 0.88 | 0.89 | 0.92 |
| OneOPES | 0.76 | 0.74 | 0.75 | 0.42 | 0.46 | 0.54 | 0.56 |
| OneOPES MultiCV | 0.74 | 0.73 | 0.73 | 0.42 | 0.44 | 0.52 | 0.55 |

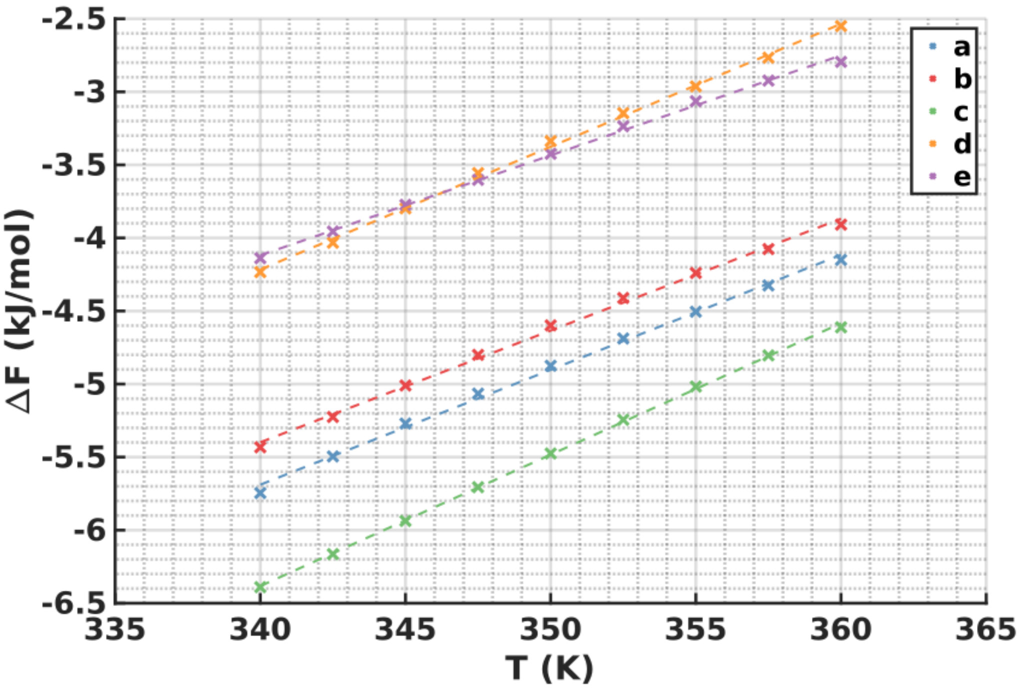


Figure S19: OneOPES MultiCV simulations of Chignolin where replica **4** is used to extract the folding $\Delta F$ in a range of temperatures. For each simulation, we perform a linear fit of $\Delta F$ against the temperature and estimate the slope and the intercept.

Table S4: Additional details about the linear fit from Fig. S19. For each simulation, we report the entropy, the enthalpy, the linear fit $R^2$, and the melting temperature.

| simulation | -$\Delta S$ (kJ/(mol·K)) | -$T\Delta S$ (kJ/(mol)) | $\Delta U$ (kJ/mol) | $R^2$ | $T_m$ (K) |
|---|---|---|---|---|---|
| a | 0.0787 | 26.7 | -32.4 | 0.997 | 412 |
| b | 0.0765 | 26.0 | -31.4 | 0.997 | 411 |
| c | 0.0900 | 30.6 | -37.0 | 0.999 | 411 |
| d | 0.0841 | 28.6 | -32.8 | 0.998 | 390 |
| e | 0.0684 | 23.3 | -27.4 | 0.998 | 400 |
